# Histone monoaminylation dynamics are regulated by a single enzyme and promote neural rhythmicity

**DOI:** 10.1101/2022.12.06.519310

**Authors:** Qingfei Zheng, Ryan M. Bastle, Shuai Zhao, Lingchun Kong, Lauren Vostal, Aarthi Ramakrishnan, Li Shen, Sasha L. Fulton, Haifeng Wang, Baichao Zhang, Akhil Upad, Lauren Dierdorff, Robert E. Thompson, Henrik Molina, Stephanie Stransky, Simone Sidoli, Tom W. Muir, Haitao Li, Yael David, Ian Maze

## Abstract

Histone H3 monoaminylations at glutamine(Q) 5 represent an important family of epigenetic markers in neurons that play critical roles in the mediation of permissive gene expression (1, 2). We previously demonstrated that H3Q5 serotonylation(ser) and dopaminylation(dop) are catalyzed by the Transglutaminase 2 (TGM2) enzyme and alter both local and global chromatin states (3, 4). Here, we found that TGM2 additionally functions as an “eraser” of H3 monoaminylations that is capable of “re-writing” these epigenetic marks in cells, including a new class of this modification, H3Q5 histaminylation(his), which displays dynamic diurnal expression in brain and contributes to neural rhythmicity. We found that H3Q5his inhibits binding of the MLL1 complex to the H3 N-terminus and attenuates its methyltransferase activity on H3 lysine(K) 4. We determined that H3Q5 monoaminylation dynamics are dictated by local monoamine concentrations, which are utilized by TGM2. Taken together, we present here a novel mechanism through which a single chromatin regulatory enzyme is capable of sensing chemical microenvironments to affect the epigenetic states of cells.

**One sentence summary:** A single enzyme, TGM2, bidirectionally controls H3 monoaminylation dynamics, which, in turn, facilitate neural rhythmicity.

## MAIN TEXT

Post-translational modifications (PTMs) of histones have emerged as a key regulatory mechanism contributing to diverse DNA-templated processes, including transcription and DNA repair (5). Well-studied PTMs, such as methylation, acetylation and ubiquitination are dynamically regulated by corresponding, site-specific “writer” and “eraser” enzymes and can be recognized by “reader” proteins that facilitate downstream cellular responses (6, 7). In addition, various small molecule metabolites can directly react with substrate proteins to form site-specific adducts, or can be indirectly added to amino acid side chains through enzymatic processes (8–10). These modifications can both impact the three-dimensional architecture of chromatin and alter transcriptional landscapes, thereby serving as important mediators of cell fate and plasticity (5–10).

We recently reported on the identification of a new class of histone PTMs, whereby monoamine neurotransmitters, such as serotonin and dopamine (termed serotonylation and dopaminylation, respectively), can be transamidated to glutamine residues (3, 4, 11–14). We specifically determined that histone H3 glutamine (Q) 5 is a primary site of modification and demonstrated that H3 monoaminylations play important roles in regulating neuronal transcriptional programs, both during development and in adult brain (3). We found that H3Q5 serotonylation (H3Q5ser) acts as a permissive mark, both enhancing the recruitment of the general transcription factor complex TFIID to the active mark H3 lysine 4 (K4) tri-methylation (me3), and attenuating demethylation of H3K4 via inhibition of KDM5 family and LSD1 demethylases (3, 12). In subsequent studies, we characterized dopaminylation events at this same site (H3Q5dop) in ventral tegmental area (VTA) and found that aberrant accumulation of H3Q5dop during abstinence to drugs of abuse promotes persistent transcriptional programs that facilitate relapse vulnerability (4, 13). Importantly, we uncovered that these monoaminylation events are catalyzed by Transglutaminase 2 (TGM2), a Ca^2+^-dependent enzyme that exhibits multiple functions, including aminolysis, hydrolysis and alcoholysis of protein glutamine residues, as well as inducing covalent protein crosslinks (15, 16). However, the precise enzymatic regulatory mechanisms through which H3 monoaminylation adducts are removed, or exchanged, to allow for neuronal transcriptional plasticity remained poorly understood.

### TGM2 is an H3 serotonylation “eraser” in cells

To investigate the regulatory mechanisms of H3 serotonylation dynamics in cells, we utilized the reported 5-propargylated tryptamine (5-PT) chemical probe to track histone serotonylation reactions via Cu(I)-catalyzed azide-alkyne cycloaddition (CuAAC) (3, 17). Initially, we performed a serotonylation pulse-chase experiment to determine the stability of H3ser in cells. We transfected HEK293T cells, which do not endogenously express TGM2, with wild-type (WT) pACEBac1-flag-TGM2 (3). Transfected cells were then treated with 500 μM 5-PT for 6 hours, washed and incubated for an additional 12 hours in 5-PT-free media. Histones were then isolated by high salt extraction, clicked with CuAAC-meditated Cyanine 5 (Cy5), separated by SDS-PAGE and imaged for fluorescence **(Figure 1A)**. Unexpectedly, we observed that the H3ser signal, which was induced in the presence of 5-PT and TGM2-WT, were decreased in TGM2-WT expressing cells during the chase phase, while histones extracted from cells treated with TGM2 inhibitors [ERW1041E (18) or ZDON (19)] did not lose the serotonylation adduct **(Figure 1B)**. These results suggested that in the absence of a serotonin donor, TGM2 may also catalyze the removal of the mark from H3. We confirmed this observation in HeLa cells, which express TGM2 endogenously, in a similar pulse-chase experiment **(Figure 1C and Supplementary Figure 1)**.

**Figure 1.**
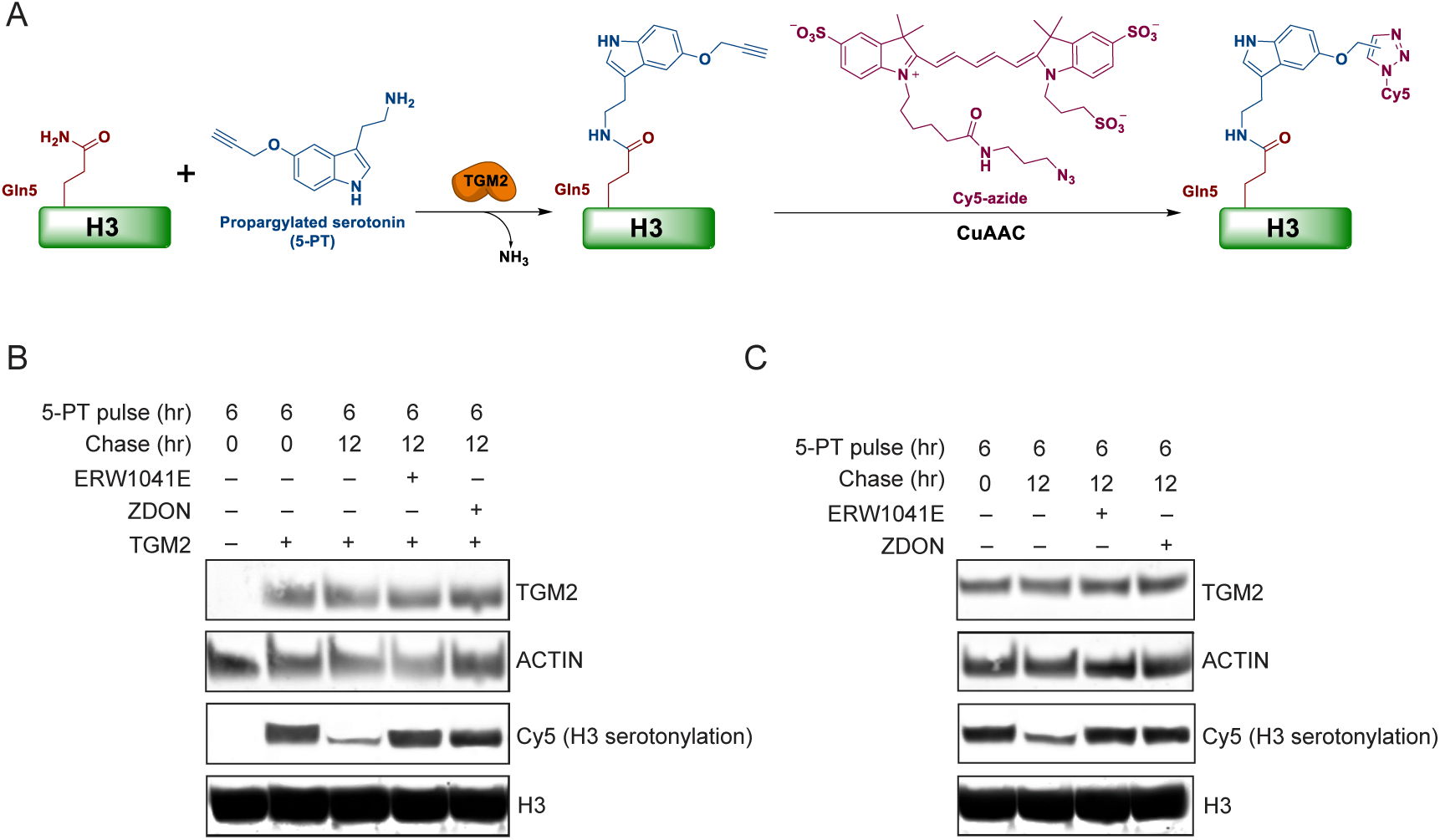
TGM2 is both necessary and sufficient for the addition and removal of H3 serotonylation *in cellulo*. (A) Schematic depicting TGM2 mediated covalent modification of H3Q5 by the serotonin analog, 5-PT, and its subsequent visualization via CuAAC-meditated Cyanine 5 (Cy5) conjugation. (B) 5-PT pulse/chase experiment in HEK293T cells, which do not endogenously express TGM2, transfected with TGM2-WT and treated -/+ TGM2 inhibitors (ERW1041E or ZDON) following the removal of 5-PT from the media. Western blotting was performed to examine Cy5 (i.e., H3 serotonylation) and TGM2 expression after media replacement -/+ TGM2 inhibition. Total H3 and ACTIN serve as loading controls. Experiment was repeated at least 3X. (C) 5-PT pulse/chase experiment in HeLa cells, which endogenously express TGM2, treated -/+ TGM2 inhibitors (ERW1041E or ZDON) following the removal of 5-PT from the media. Western blotting was performed to examine Cy5 (i.e., H3 serotonylation) and TGM2 expression after media replacement -/+ TGM2 inhibition. Total H3 and ACTIN serve as loading controls. Experiment was repeated at least 3X.

### TGM2 is an H3 monoaminylation “re-writer” in vitro

Since our results suggested a serotonylation erasing capability for TGM2, we aimed to test whether this mechanism relies on its canonical catalytic residue, cysteine (Cys) 277, through a nucleophilic attack of the γ-carboxamide group of glutamines **(Figure 2A)**. In this hypothesized mechanism, a thioester intermediate would then activate the amide bond to accept a second nucleophilic attack from donors within the reaction’s microenvironment. To test this, we first synthesized H3 N-terminal tail peptides (1-21 amino acids) to act as substrates and product standards for Q5ser, Q5dop and Q5E (the predicted deamidated product) **(Supplementary Figure 2)**. Next, H3Q5ser or H3Q5dop peptides were incubated with lysates from HEK293T cells expressing either TGM2-WT or the catalytically dead mutant, TGM2-C277A. Liquid chromatography-mass spectrometry (LC-MS) analyses revealed that H3Q5ser and H3Q5dop peptides incubated with lysates containing TGM2-WT underwent stoichiometric removal/deamidation [see **Figure 2B, xi** for LC-MS trace of the deamidated product – Q5 to E (glutamic acid) 5] of the serotonylation and dopaminylation adducts. Importantly, the monoaminylated peptides remained unchanged in the presence of lysates from cells expressing the mutant TGM2-C277A enzyme **(Figure 2B, i-iv)**. To provide more insights into the possible biochemical mechanism of TGM2-dependent monoaminylation “re-writing” activity, we purified recombinant TGM2-WT and TGM2-C277A to be used in *in vitro* reactions **(Supplementary Figure 3)**. Next, we incubated either WT or mutant TGM2 enzymes with H3Q5ser or H3Q5dop peptides in the presence of Ca^2+^. We similarly analyzed these *in vitro* reactions by LC-MS, which provided consistent results to those observed after cell lysate incubations **(Figure 2B, v-viii)**. Co-immunoprecipitation (Co-IP) experiments performed on recombinant H3 proteins confirmed this unique enzymatic mechanism by capturing the putative TGM2-H3 thioester complex with TGM2-WT but not with TGM2-C277A **(Supplementary Figure 4)**.

**Figure 2.**
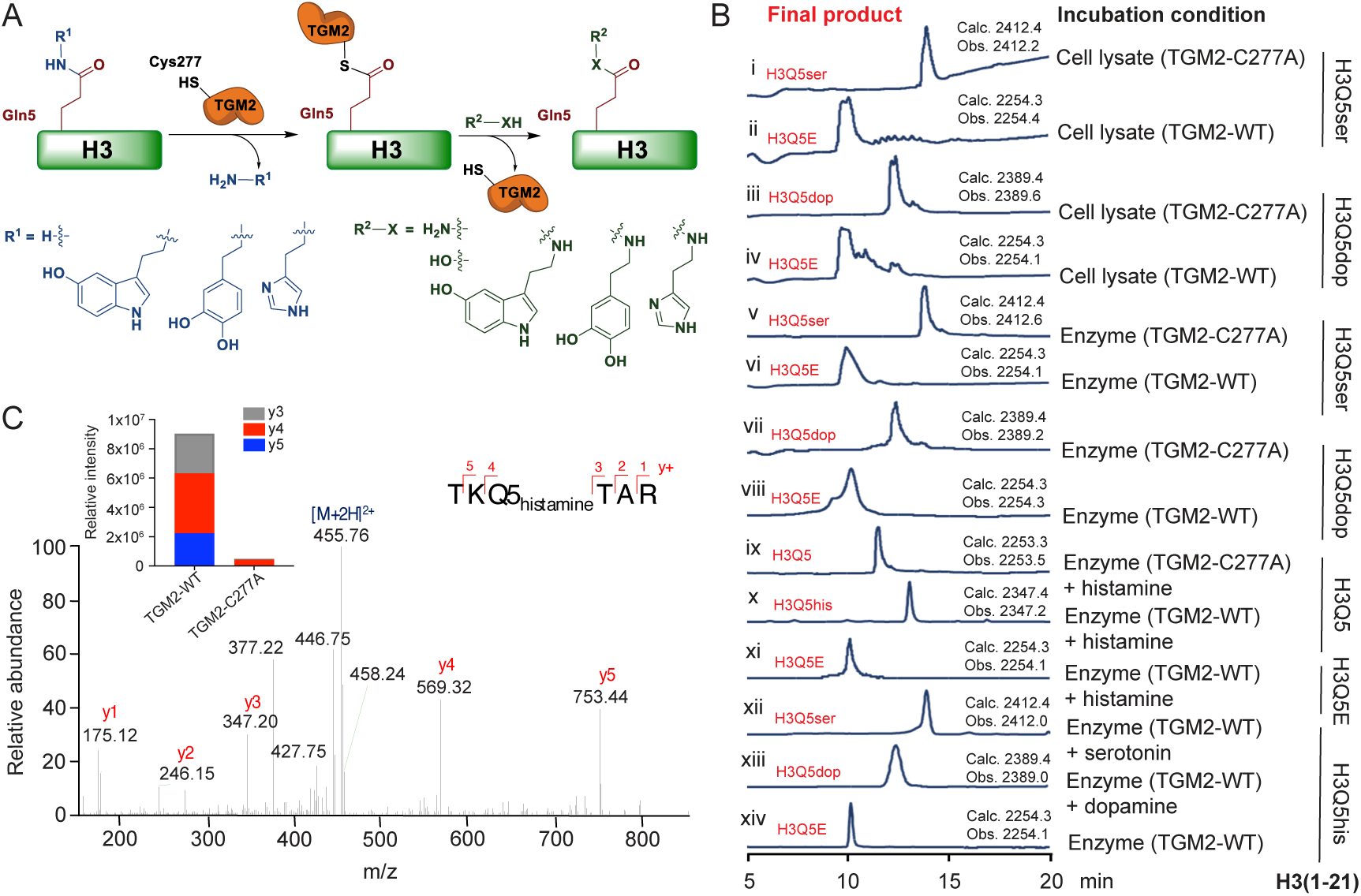
TGM2 “writes,” “erases” and “re-writes” H3 monoaminylations *in vitro*. (A) Hypothesized mechanism of TGM2 mediated H3 monoaminylation ‘writing,’ ‘erasing’ and ‘re-writing.’ In this model, TGM2’s catalytic residue, cysteine (Cys) 277, attacks the γ-carboxamide group of glutamine 5 on H3, and the formation of a thioester intermediate structure then activates the amide bond to accept a nucleophilic attack from donors within the local microenvironment. (B) LC-MS analyses of synthetically modified H3 (1–21) peptides – H3Q5ser *vs.* H3Q5dop *vs.* H3Q5his (as well as an H3Q5E peptide standard for assessments of H3Q5 deamidation; xi) – following incubation with either cellular lysates expressing TGM2-WT or TGM2-C277A (i-iv) or recombinant TGM2-WT *vs.* TGM2-C277A (v-x). H3Q5his peptides were further incubated with TGM2-WT in the presence or absence of a replacement monoamine donor (xii – serotonin *vs.* xiii – dopamine), demonstrating that TGM2-WT can both deamidate H3 monoaminylations in the absence of a replacement donor (xiv) and ‘re-write’ H3 monoaminylations in the presence of replacement donors. Calculated *vs.*observed masses are provided. HPLC UV traces, λ = 214 nm. Experiment was repeated at least 3X. (C) LC-MS/MS analyses identified endogenous H3Q5his in HEK293T cells transfected with TGM2-WT following histamine treatments. y+ ions are annotated. Insert: Relative intensities of the most abundant *y* fragment ions from the H3Q5his peptide.

Given the “re-writing” activity of TGM2 and the physiological significance of histamine, another monoamine molecule involved in diverse physiological processes ranging from local immune signaling to functions of the gut and neurotransmission (20), we next focused our attention on histamine as a potential alternative donor for histone H3Q5 monoaminylation (H3Q5his). To test whether histamine, like serotonin and dopamine, can act as a cofactor for TGM2 to modify H3Q5, we examined this reaction in a purified system. We incubated WT H3 peptides with recombinant TGM2 (WT or C277A) in the presence of histamine, followed by LC-MS analysis **(Figure 2B, ix-x and Supplementary Figure 5A-D)**. Our results indicated that histamine was stoichiometrically added to the H3 peptide at glutamine 5 (**Supplementary Figure 5C**), consistent with previous reports suggesting that histamine is a preferred donor for TGM2 mediated transamidation of other substrate proteins (21). Moreover, we found that the synthetic H3Q5his peptide was converted to H3Q5E by TGM2-WT in the absence of free histamine **(Figure 2B, xiv and Supplementary Figure 2)** and was fully converted to H3Q5ser **(Figure 2B, xii)** or H3Q5dop **(Figure 2B, xiii)** in the presence of the corresponding monoamine donor. Importantly, we identified the presence of endogenous H3Q5his in HEK293T cells expressing TGM2-WT and treated with histamine, but not in cells expressing TGM2-C277A, via LC-MS/MS **(Figure 2C)**.

### TGM2 regulates H3Q5his dynamics in vitro and in cells

To identify whether H3Q5his accumulates in cells when histamine is provided as a donor, we generated a polyclonal antibody that selectively recognizes H3Q5his, but not the other monoaminyl marks, *in vitro* and *in vivo* (using cells and brain tissues) – see **Supplementary Figure 6A-E** for antibody validations. We treated HEK293T cells expressing TGM2-WT or TGM2-C277A with histamine, and following cell harvesting, histones were extracted and analyzed by western blot analysis using the new antibody. These results revealed the accumulation of H3Q5his in TGM2-WT expressing cells (*vide infra,* **Figure 3B**).

**Figure 3.**
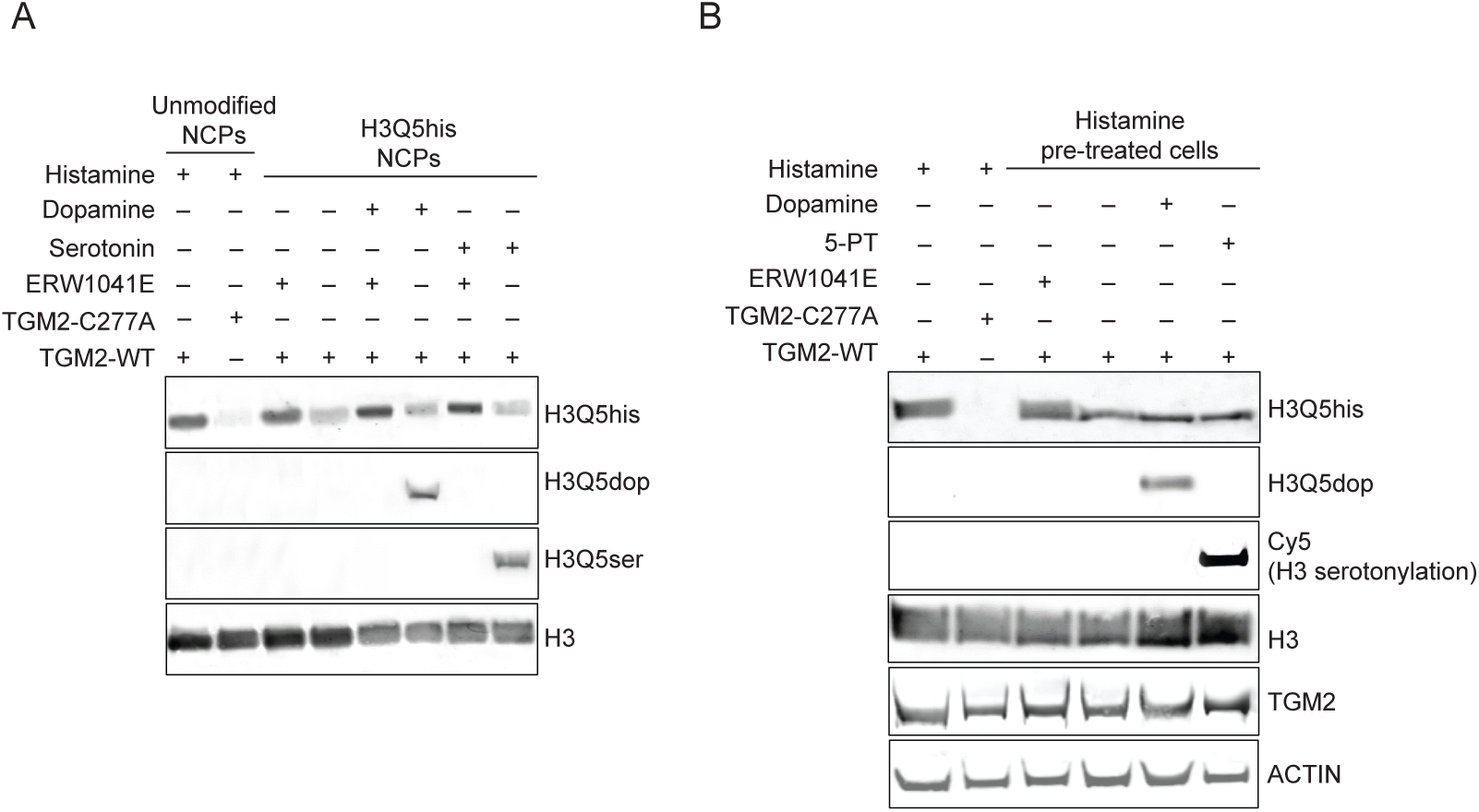
Noncanonical TGM2 mediated “re-writing” of H3 monoaminylations on NCPs and in cells. (A) TGM2-WT, but not TGM2-C277A, can transamidate histamine to H3Q5 on NCPs. NCPs pre-modified by histamine at H3Q5 can be deamidated by TGM2-WT, a mechanism that is inhibited by treatment with the TGM2 inhibitor ERW1041E. TGM2-WT can additionally ‘re-write’ H3Q5his on NCPs in the presence of replacement monoamines donors, such as serotonin or dopamine, resulting in the establishment of H3Q5ser or H3Q5dop, respectively. Experiment repeated at least 3X. (B) TGM2-WT, but not TGM2-C277A, can transamidate histamine to H3Q5 in HEK293T cells. H3Q5his pre-modified histones can be deamidated by TGM2-WT *in cellulo*, a mechanism that is inhibited by treatment with the TGM2 inhibitor ERW1041E. TGM2-WT can also ‘re-write’ H3Q5his *in cellulo* in the presence of replacement monoamines donors, such as 5-PT (serotonin analog; assessed via Cy5 conjugation) or dopamine, resulting in the establishment of H3 serotonylation or H3Q5dop, respectively. Total H3 and ACTIN western blots are provided as loading controls. Experiment repeated at least 3X.

To further investigate the dynamics of H3Q5his and its regulation by TGM2 in a more physiologically relevant context, we performed an *in vitro* competition assay using reconstituted nucleosome core particles (NCPs) as substrates **(Figure 3A)**. We performed pulse-chase experiments, where we treated NCPs with one monoamine (in this case histamine) and used a buffer exchange column to replace the monoamine donor before the chase. Our results not only indicated that H3Q5his can be efficiently removed from NCPs in a TGM2-dependent manner, but also that the presence of a different monoamine donor during the chase phase resulted in “re-writing” of the mark to alternative monoaminylation states based upon the donor provided. This reaction did not occur in the presence of the TGM2-specific inhibitor, ERW1041E **(Figure 3A)**.

Next, a similar pulse-chase experiment was performed using HEK293T cells expressing TGM2. Histamine was added to the media to allow for H3Q5his establishment, followed by a chase with media containing 5-PT or dopamine. The analysis of H3 monoaminylation, presented in **Figure 3B**, demonstrated that in the presence of TGM2, H3Q5his was efficiently converted to the alternative monoaminylation state as a function of the donor. Overall, these results indicated that histone monoaminylation is a family of highly dynamic epigenetic modifications that are regulated by a single enzyme, TGM2, based upon the local concentration of monoamine donors within a given cellular context.

### H3Q5 histaminylation is enriched in tuberomammillary nucleus and displays dirurnal rhythmicity

To explore the biological significance of H3Q5his dynamics *in vivo*, we turned our investigations to the posterior hypothalamic tuberomammillary nucleus (TMN), which consists largely of histaminergic neurons and is involved in diverse biological functions ranging from control of arousal to maintenance of sleep-wakes cycles and energy balance (22). Indeed, TMN was found to be enriched for H3Q5his *vs.* other non-histaminergic nuclei examined **(Supplementary Figure 6I)**. Given the TMN’s prominent role in diurnal behavioral rhythmicity, we first assessed whether the TMN itself displays rhythmic patterns of gene expression that may require chromatin-based control. To do so, we performed bulk tissue RNA-seq on mouse TMN tissues collected at various time points across the zeitgeber (ZT) beginning at time ZT0 hour (the beginning of ‘lights on’ for mice, which marks the initiation of their inactive phase owing to nocturnalism) and then every 4 hours for 24 hours, with ZT12 marking the beginning of the animals active phase. These RNA-seq data were then analyzed using JTK_CYCLE (23), which is a non-parametric algorithm designed to detect rhythmic components in genome-wide datasets. These analyses revealed that the TMN displays rhythmic gene expression patterns **(Figure 4A)**, with many known circadian genes (e.g., *Arntl*, *Dbp*, *Per1/2*, etc.) identified as being significantly regulated in this manner **(Figure 4B and Supplementary Data 1)**. Furthermore, ChEA analysis (24), which infers transcription factor regulation from integration of previous genome-wide chromatin immunoprecipitation (ChIP) analyses, identified CLOCK as a robust transcriptional regulator of the genes identified in our rhythmic gene expression dataset **(Figure 4C)**, thereby confirming the histaminergic TMN as a transcriptionally rhythmic brain region.

**Figure 4.**
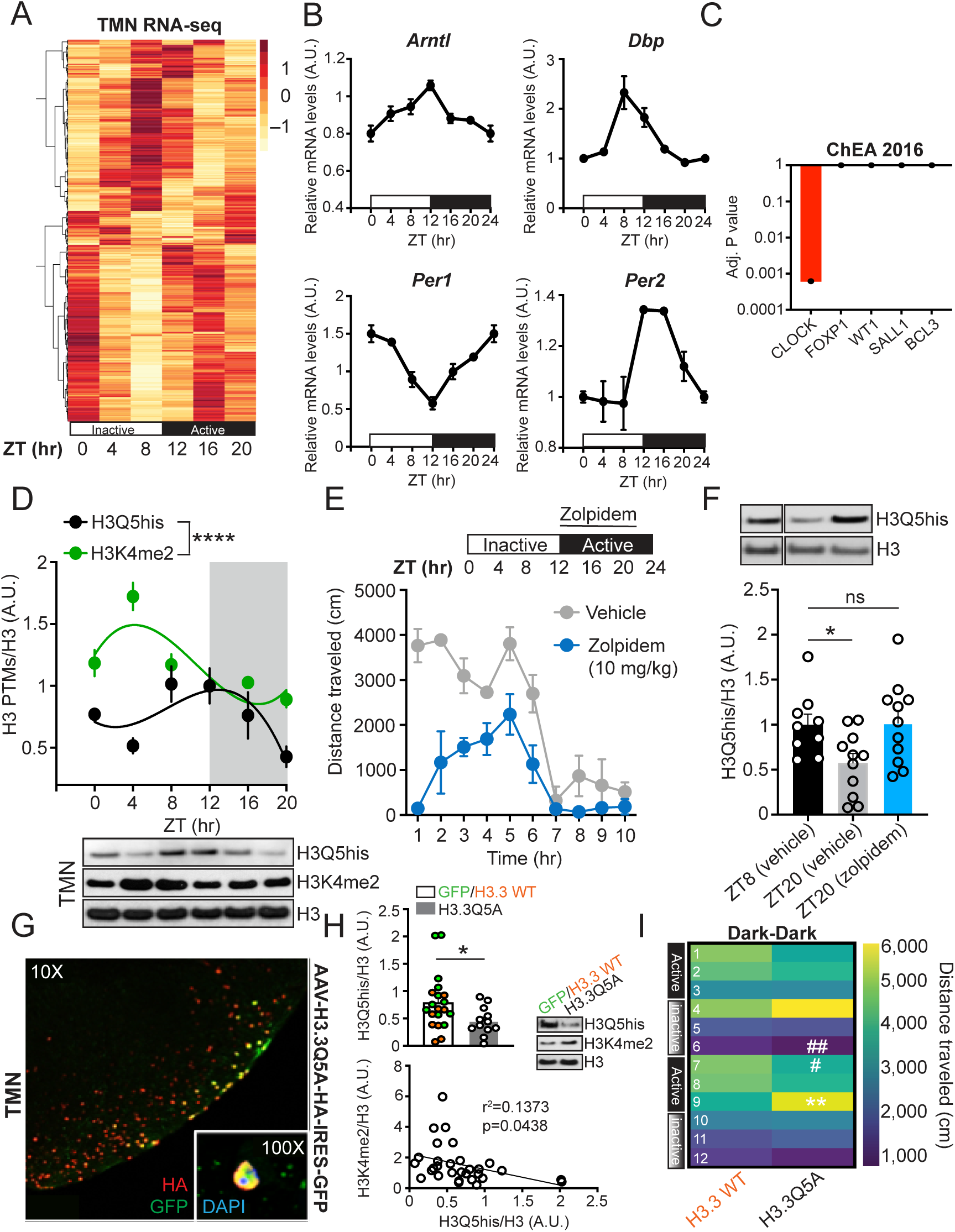
Diurnal H3Q5his dynamics in TMN contribute to neural rhythmicity. (A) RNA-seq of mouse TMN across the ZT (tissues collected at ZT 0, 4, 8, 12, 16 and 20) identifies rhythmic patterns of gene expression (*n*=3 biological replicates/time point), (B) including known circadian genes, such as *Arntl*, *Dbp*, *Per1* and *Per2*. (C) ChEA analysis reveals the transcription factor CLOCK as an important contributor to rhythmic genes identified in TMN. (D) H3Q5his in TMN displays a rhythmic pattern of expression across the ZT in mice [*n*=8-10 per time point; comparison of fits analysis between third order polynomial, cubic trend (alternative hypothesis), *vs.* first order polynomial, straight line (null hypothesis) – p=0.0034, null hypothesis rejected], a pattern that significantly diverges from that of adjacent H3K4me2 (p<0.0001), a mark that also displays rhythmic expression in this brain region (*n*=6-10 per time point; comparison of fits between third order polynomial, cubic trend (alternative hypothesis), *vs.* first order polynomial, straight line (null hypothesis) – p=0.0079, null hypothesis rejected]. H3 western blots are provided as loading control. (E) Assessments of locomotor activity in mice treated with Zolpidem *vs.* vehicle during their active phase (*n*=10/group). (F) Examination of H3Q5his in TMN of mice treated with Zolpidem *vs.* vehicle during their active phase at ZT8 (vehicle, *n*=9) and ZT 20 (Zolpidem *vs.* vehicle, (*n*=11/group) – one-way ANOVA (p=0.0227, Dunnett’s post hoc test, *p<0.05. H3 western blots are provided as loading control. (G) IHC/IF analysis confirming nuclear expression of H3.3Q5A-HA dominant negative in TMN of mice expressing AAV-H3.3Q5A-HA-IRES-GFP. (H) Top – Western blot validation of H3Q5his downregulation in TMN following AAV mediated expression of H3.3Q5A *vs.* H3.3WT or empty vector controls (*n*=9-12/viral treatment group – control groups collapsed; unpaired student’s t-test, *p=0.0422). Bottom – Linear regression analysis comparing intra-animal H3Q5his *vs.* H3K4me2 expression in TMN following viral infections with vectors expression H3.3Q5A *vs.* H3.3 WT or empty vector controls (*n*=9-12/viral treatment group). H3 western blots are provided as loading control for both marks. (I) Following intra-TMN transduction with H3.3Q5A *vs.* H3.3 WT, mice were monitored for locomotor activity beginning 12 hours after a shift from light-dark to dark-dark to examine whether disrupting H3Q5his alters normal circadian cycling activities. Disrupting normal H3Q5his dynamics in TMN resulted in abnormal shifts in diurnal locomotor activity during transitions from inactive to active states, and vice versa. Heatmap presents locomotor data binned into 4-hour intervals for a total of 48 hours [*n*=10-11/viral group, two-way ANOVA, interaction of time x virus – **p=0.0040, Sidak’s multiple comparison test – **p=0.0079, *a posteriori* Student’s t-test – #p<0.05, ##p<0.01]. Data are represented as mean ± SEM.

We next performed western blotting for H3Q5his in TMN across the ZT to determine the mark’s expression pattern. We found that H3Q5his is a rhythmic histone modification, with its expression highest during transitions into the active phase and lowest during transitions into the inactive phase for mice **(Figure 4D)**. Such patterns of expression appeared to mimic those of the *Hdc* gene **(Supplementary Figure 7A)** that encodes for the protein Histidine decarboxylase, which mediates the catalysis of histidine to form histamine, suggesting that fluctuations in histamine metabolism may contribute to overall levels of the mark in this brain region.

Given that our previous data indicated that other H3 monoaminylation states often function combinatorially with adjacent H3K4 methylation (specifically H3K4me3) to control transcriptional processes (8–9), we next generated a site-specific antibody that selectively recognizes H3K4me3Q5his (see **Supplementary Figure 6F-H** for antibody validations), followed by expression profiling across the ZT in TMN. We found that H3K4me3Q5his, while present in TMN, does not display a rhythmic pattern of expression, with similar results observed using an H3K4me3 specific antibody, which recognizes H3K4me3 both in the absence and presence of Q5his **(Supplementary Figure 7B)**. H3K4me1 and TGM2 were also found to be non-rhythmic **(Supplementary Figure 7B)**, providing further evidence that H3Q5his dynamics are a function of histamine availability rather than TGM2 expression. While our data indicated that H3K4me1, H3K4me3 and H3K4me3Q5his were not rhythmic in TMN, previous literature had demonstrated that other valence states of H3K4 methylation, such as H3K4me2, may also contribute to diurnal gene expression (25). As such, we examined H3K4me2 expression across the ZT, and found that not only is H3K4me2 rhythmic but also that its pattern of expression significantly diverges from that of H3Q5his, suggesting that H3Q5his may play an antagonistic role in the regulation of H3K4me2 dynamics **(Figure 4D)**. Finally, to assess whether other brain regions that receive histaminergic projections also display H3Q5his dynamics related to diurnal cycling, we evaluated H3Q5his expression across the ZT in the hypothalamic suprachiasmatic nucleus (SCN), which is considered to be a brain region important for producing circadian rhythms (26). Unlike in TMN, H3Q5his was not found to be rhythmic in SCN **(Supplementary Figure 7C)**.

### Diurnal fluctuations of H3Q5his in TMN contribute to molecular and behavioral rhythmicity

We next sought to investigate whether perturbing diurnal rhythms may result in altered regulation of H3Q5his dynamics in TMN, and reciprocally, whether altering H3Q5his may disrupt diurnal gene expression and behavior. To do so, we treated mice pharmacologically with the sleep aid Zolpidem (aka Ambien) at 10 mg/kg during the beginning of the mouse active phase – a perturbation that robustly results in an immediate loss of activity, as measured by locomotor behavior **(Figure 4E).** We then collected TMN tissues 8 hours after treatment for assessments of H3Q5his expression. In vehicle treated controls, comparing H3Q5his expression at ZT8 *vs.* ZT20, we found that H3Q5his expression is reduced, as expected. However, treatment with Zolpidem significantly attenuated this reduction in expression **(Figure 4F)**, suggesting that the mark’s levels are linked to sleep-wake cycles. While H3Q5his expression can be altered by inducing sleep, it remained unclear whether the mark plays a causal role in molecular or behavioral rhythmicity. To investigate this, we next performed Adeno-associated viral vector (AAV) transduction in TMN to introduce a mutant H3, H3.3Q5A, which is actively incorporated into neuronal chromatin **(Figure 4G)** and functions as a dominant negative, thereby reducing global levels of the mark (H3.3Q5A) (11). In these experiments, two AAV controls were transduced: an empty vector control expressing GFP and H3.3 WT, which were collapsed as a common control since H3.3 WT transduction did not alter H3Q5his expression *vs.* GFP **(Figure 4H, top)**. We additionally assessed H3K4me2 in response to viral transduction and found that its expression was anti-correlated with H3Q5his **(Figure 4H, bottom)**, consistent with the rhythmic expression data presented in **Figure 4D**. Finally, we examined what impact such perturbations may have on diurnal behavior, as assessed via locomotor activity across the ZT. Three weeks following viral transductions (the time required to achieve maximal expression of the viruses), mice were monitored for locomotor activity beginning 12 hours after a shift from light-dark (i.e., their normal housing conditions) to dark-dark in order to examine whether disrupting H3Q5his alters normal circadian cycling activities. Consistent with our hypothesis, we found that perturbing normal H3Q5his dynamics in TMN resulted in abnormal shifts in diurnal locomotor activity, particularly during transitions from inactive to active states, and vice versa **(Figure 4I)**. Subsequent RNA-seq on transduced TMN tissues confirmed significant overlaps with genes disrupted by H3Q5his manipulations in comparisons to those identified as being rhythmic in **Figure 1A-C**, as well as those predicted to be CLOCK gene targets **(Supplementary Figure 7D and Supplementary Data 2)**. In sum, our data revealed that not only is H3Q5his dynamic in its expression across the ZT but also that it contributes significantly to rhythmic gene expression and behavior.

### H3Q5his antagonizes H3K4 methyltransferase activity through electrostatics

To investigate the potential biochemical mechanism governing H3Q5his’ effect on H3K4 methylation dynamics during diurnal cycling – and, in turn, rhythmic gene expression and behavior – we next sought to explore whether H3Q5his may directly alter the activity of the H3K4 methylation machinery *in vitro*. Given that previous work, including our own, indicated that H3Q5ser can potentiate binding to WDR5 (3, 12, 27), an H3 “reader” protein and member of the MLL1 methyltransferase complex (28), we began by performing MALDI-TOF mass spectrometry on both unmodified H3 and H3Q5his peptides (1–15). We found that H3Q5his robustly attenuated MLL1 complex processivity to add methylation to H3K4 **(Figure 5A)**. These results were orthogonally confirmed using LC-MS/MS to monitor the establishment of H3K4 methylation states by MLL1 on H3 (1–21) unmodified *vs.* Q5ser *vs.* Q5his peptides, where we found that while H3Q5ser does not alter MLL1 activity *vs.* unmodified H3, H3Q5his significantly attenuates MLL1 complex activity (**Figure 5B-C**).

**Figure 5.**
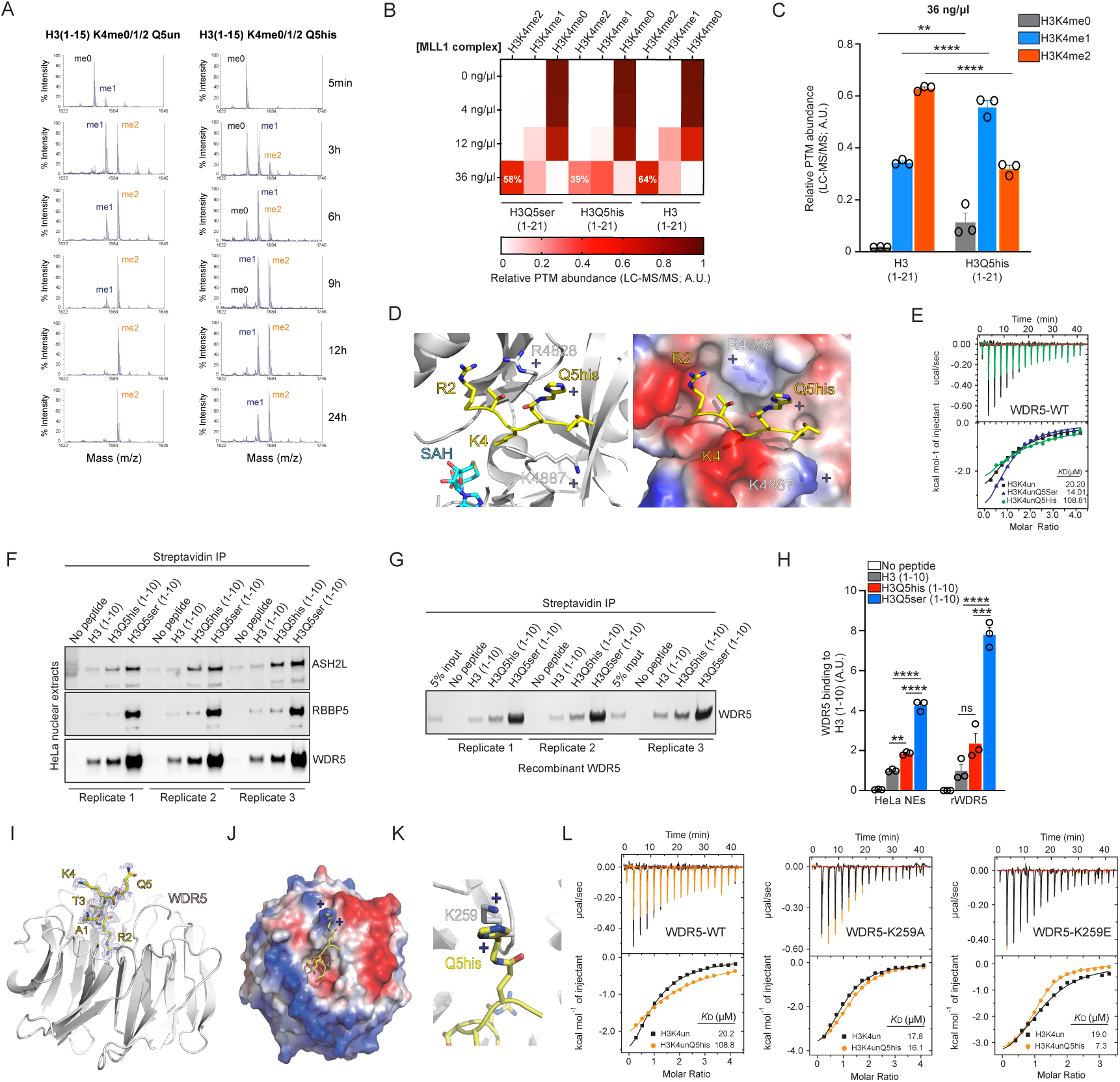
H3Q5his antagonizes MLL1 mediated H3K4 methylation establishment via electrostatics. (A) MALDI-TOF assessments of the recombinant MLL1 complex methyltransferase activities on H3K4 comparing H3K4unmodQ5unmod *vs.* H3K4unmodQ5his peptides (1–15). (B) LC-MS/MS quantification of H3K4 methylation states (H3K4me0 *vs.* H3K4me1 *vs.* H3K4me2; H3K4me3 signal was negligible and was thus omitted) on H3 (1–21) unmodified *vs.* H3Q5ser *vs.* H3Q5his peptides, titrating the concentration of MLL1 complex in the system. (C) LC-MS/MS quantification of H3K4 methylation states (H3K4me0 *vs.* H3K4me1 *vs.* H3K4me2; H3K4me3 signal was negligible and was thus omitted) on H3 (1–21) unmodified *vs.* H3Q5ser *vs.* H3Q5his peptides using 36 ng/ul of MLL1 complex with replicates (*n* = 3) for quantification. Two-way ANOVA with Sidak’s multiple comparison test; **p<0.01, ****p<0.0001. (D) The catalytic center of MLL3 (PDB ID: 5F6K). The H3Q5his modification was modeled into the structure. (E) ITC assessments of WDR5_WD40_ binding to H3K4unmodQ5unmod *vs.* H3K4unmodQ5his *vs.* H3K4unmodQ5ser peptides. (E) ITC assessments of WDR5_WD40_ binding to H3K4unmodQ5unmod *vs.* H3K4unmodQ5his *vs.* H3K4unmodQ5ser peptides. (F) Peptide IPs [H3 (1–10) unmodified *vs.* Q5his *vs.* Q5ser] against soluble nuclear extracts from HeLa cells comparing their interactions with members of the MLL1 complex (ASH2L, RBBP5 and WDR5) via western blotting. (G) Results from (F) were orthogonally confirmed by peptide IPs against recombinant full length WDR5. (H) Quantification of binding data from (F-G). Within experiment one-way ANOVAs followed by Tukey’s multiple comparison tests; **p<0.01, ***p<0.001, ****p<0.0001. (I) X-ray crystal structure of the WDR5_WD40_-H3 complex demonstrated that H3Q5 displays flexible surface binding with WDR5_WD40_. (J-K) The location of Q5 near the positively charged WDR5_WD40_K259 residue in the structure indicated that WDR5_WD40_K259 and H3Q5his may serve to electrostatically repel one another, resulting in binding inhibition. (L) Left – ITC assessments of WDR5_WD40_ binding to H3K4unmodQ5unmod *vs.* H3K4unmodQ5his peptides. Middle – ITC assessments of WDR5_WD40_K259A binding to H3K4unmodQ5unmod *vs.* H3K4unmodQ5his peptides. Right – ITC assessments of WDR5_WD40_K259E binding to H3K4unmodQ5unmod *vs.* H3K4unmodQ5his peptides.

Based upon this attenuation, we next wished to explore the potential mechanisms responsible for this phenomenon. We began by modeling possible H3Q5his interactions with MLL using the published MLL3-RBBP5-ASH2L-H3 complex structure (29). Based upon our structural modeling assessments (**Figure 5D**), we predict that the H3Q5 residue would be ‘sandwiched’ between arginine (R) 4828 and K4887 of MLL3, and while enough space would exist, in theory, to tolerate a large modification, such as serotonylation (with stacking between R4828 and serotonylation possibly stabilizing the H3 peptide; in addition, the double conformation of WDR5 F149 supports the notion that the binding surface of H3Q5ser is flexible, **Supplementary Figure 8**). However, the positively charged histaminylation moiety would not be favored owing to electrostatics. Additionally, since the MLL1 complex member WDR5 (which presents the H3 tail to MLL1 for deposition of H3K4 methylation) has previously been shown to interact with H3Q5 serotonylation through its WD repeats (12, 27), we next wished to assess whether H3Q5his may antagonize such interactions *vs.* H3Q5ser. We performed isothermal titration calorimetry (ITC) to measure interactions between the purified WD40 domain of WDR5 (WDR5_WD40_) and the monoaminylated H3 tail (H3Q5ser *vs.* H3Q5his). As expected, WDR5_WD40_ was found to bind to the unmodified H3 tail (K_D_ 20.20 μM), with a slight increase in binding observed in the context of a H3Q5ser substrate (K_D_ 14.01 μM). Interestingly, H3Q5his was observed to attenuate this binding (K_D_ 108.81 μM), reducing the affinity between H3 and WDR5_WD40_ by ∼5-fold **(Figure 5E**; see **Supplementary Table 1 for ITC statistics)**.

Since the MLL1 complex binds the H3 tail directly via interactions with WDR5, which can facilitate MLL1 methyltransferase activities (30), we next performed peptide IPs [H3 (1–10) unmodified *vs.* Q5his *vs.* Q5ser] against soluble nuclear extracts from HeLa cells. This was used to compare the interactions between the H3 peptides and members of the MLL1 complex (ASH2L, RBBP5 and WDR5) via western blotting. We found that while H3Q5ser significantly potentiates interactions between the MLL1 complex and the H3 tail, H3Q5his displayed abrogated binding (**Figure 5F** and quantified in **Figure 5H, left**). These results were orthogonally confirmed by performing peptide IPs against recombinant full length WDR5 (**Figure 5G** and quantified in **Figure 5H, right**). Given that H3Q5his, unlike H3Q5ser, is positively charged at physiological pH, we next explored whether its charge may contribute to the inhibition of binding *vs.* H3Q5ser. To this end, we used X-ray crystallography to determine the structure of the WDR5_WD40_-H3 complex in the presence of an H3Q5his peptide. While the electron density of H3Q5 was traceable within the structure, the density around the histamine moiety was not resolved **(Figure 5I**, and **Supplementary Table 2 for X-ray crystallography data collection and refinement statistics)**. We were, however, able to effectively model H3Q5his interactions based upon the orientation of the H3Q5 residue, restricted within the Ramachandran plot. This modeling indicated that while the H3R2 sidechain was inserted into the binding pocket of WDR5_WD40_ (with the H3K4 sidechain at the surface of WDR5_WD40_), as was shown previously (31), H3Q5his was predicted to display surface binding adjacent to the binding pocket. Additionally, the location of Q5 in proximity to the positively charged WDR5_WD40_K259 residue in the structure suggested that WDR5_WD40_K259 and H3Q5his may serve to electrostatically repel one another, thereby resulting in binding inhibition **(Figure 5J-K)**. To test this directly, we performed additional ITC analyses using WDR5_WD40_ mutants. We found that while H3Q5his inhibits WDR5_WD40_ binding to the H3 tail, WDR5_WD40_K259A rescued this interaction **(Figure 5L;** see **Supplementary Table 1 for ITC statistics)**. Reciprocally, the WDR5_WD40_K259E mutant displayed a slight increase in binding of WDR5_WD40_ to H3Q5his.

In summary, we investigated the dynamics of monoamine installation by TGM2, which revealed that that these PTMs unexpectedly and directly compete. Mechanistically, this is due to the capacity of TGM2 to act as “writer,” “eraser” and “re-writer” of the monoaminylated moieties on H3. In fact, we found that TGM2 can utilize any of the monoamine cofactors examined on either unmodified or monoaminylated glutamines, a process that is primarily dependent on the microenvironmental concentrations of provided nucleophiles. Furthermore, when TGM2 “erases” the monoamine adduct in the absence of alternative donors, it leaves a glutamate at the site of modification, which results in a form of “protein-induced mutagenesis” that would likely prove deleterious to cellular function. As such, TGM2’s “re-writing” activity on H3 likely represents a critical mechanism for maintaining cellular homeostasis during periods of intracellular monoamine fluctuations.

Leveraging this mechanistic insight, we identified a third class of H3 monoaminylation, histaminylation, which is also facilitated by TGM2 and occurs at the same site on H3 (Q5). Given previous evidence indicating that histamine levels in neonatal rodent brain are largely nuclear, the functions of which have remained elusive, we were interested in further exploring mechanistic roles for this new PTM *in vivo* (32). Using a new site-specific antibody that we developed, we found that H3Q5his expression is enriched at the site of histamine production in brain and showed that H3Q5his expression is dynamic in TMN as a function of sleep-wake cycles. These opposed patterns of H3K4 di-methylation, which have previously been implicated in circadian gene expression (25). Furthermore, we found that such dynamics can be disrupted by pharmacological manipulations of sleep behavior. In support, intra-TMN viral manipulations that reduce H3Q5his expression were found to disrupt normal patterns of locomotor activity during sleep-wake transitions, suggesting that the mark’s dynamics may contribute to diurnal related phenotypes. Finally, based upon a series of biochemical and structural assessments, we found that H3Q5his, in comparison to H3Q5ser, attenuates MLL1 complex binding to the H3 tail. This is specifically governed by a predicted electrostatic repulsion (owing to H3Q5his’ positive charge), and inhibits MLL1 complex mediated establishment of H3K4 methylation. Taken together, these data provide a previously unreported paradigm that addresses the dynamic regulation of histone monoaminylation events in cells/brain. Future studies aimed at elucidating TGM2’s “re-writing” capabilities in brain (e.g., switching between H3 monoaminylation states within neural cell-type specific contexts to mediate chromatin state transitions that influence transcription) are warranted and may explain, at least in part, how monoamine dynamics – both adaptive and maladaptive – contribute to brain plasticity.

## ACKNOWLEDGEMENTS

We would like to thank members of the Maze and David laboratories for critical readings of the manuscript. This work was partially supported by grants from the National Institutes of Health: R01 MH116900 (I.M.), R35 GM138386 (Y.D.), P30 CA008748 (Y.D.), P50 CA192937 (Y.D.), T32 DA007135 (R.M.B.), S10 OD030286 (S.S.), P30 CA013330 (S.S.), R37 GM086868 (T.W.M.), as well as funds from OSUCCC (Q.Z.), AFAR (Sagol Network GerOmics award; S.S.), Deerfield (Xseed award; S.S.), Relay Therapeutics (S.S.), Merck (S.S.), and the Einstein-Mount Sinai Diabetes Research Center (S.S.). The David Lab is also supported by the Josie Robertson Foundation, the Pershing Square Sohn Cancer Research Alliance, the Parker Institute for Cancer Immunotherapy, the STARR Cancer Alliance award, and the Anna Fuller Trust. In addition, the David lab is supported by W. H. Goodwin, A. Goodwin, and the Commonwealth Foundation for Cancer Research and the Center for Experimental Therapeutics at MSKCC.

## Contributions

I.M., Y.D. and H.L. conceived of the project with input from Q.Z., R.M.B, S.Z., & T.W.M. Q.Z., R.M.B., S.Z., S.S., H.L., Y.D., & I.M. designed the experiments and interpreted the data. Q.Z., R.M.B., S.Z., L.K., L.V., H.F., B.Z., A.U., L.D., R.E.T., H.M., S.S., S.S., T.W.M., H.L., Y.D. & I.M. collected and analyzed the data. A.R. & L.S. performed the sequencing-based bioinformatics with input from S.L.F. Q.Z., R.M.B., H.L., Y.D. & I.M. wrote the manuscript with input from other authors.

## Data availability

The RNA-seq data generated in this study have been deposited in the National Center for Biotechnology Information Gene Expression Omnibus (GEO) database under accession number GSE209834. Raw files for the H3Q5his mass spectrometry proteomics data in 293T cells have been deposited to the Chorus repository under project #1782. The atomic coordinates and structure factors have been deposited in the Protein Data Bank (PDB) under PDB ID code 8HMX. We declare that the data supporting findings for this study are available within the article and Supplementary Information. Related data are available from the corresponding author upon reasonable request. No restrictions on data availability apply.

## Competing interests

The Authors declare no competing interests

## MATERIALS AND METHODS

### General Methods (Equipment, Reagents, Chemicals)

UV spectrometry was performed on a NanoDrop 2000c (Thermo Scientific). Biochemicals and media were purchased from Fisher Scientific or Sigma-Aldrich Corporation unless otherwise stated. T4 DNA ligase, DNA polymerase and restriction enzymes were obtained from New England BioLabs. PCR amplifications were performed on an Applied Biosystems Veriti Thermal Cycler using either Taq DNA polymerase (Vazyme Biotech) for routine genotype verification or Phanta Max Super-Fidelity DNA Polymerase (Vazyme Biotech) for high-fidelity amplification. Site-specific mutagenesis was performed according to standard procedures of the QuickChange Site-Directed Mutagenesis Kit purchased from Stratagene (GE Healthcare) or Mut Express II (Vazyme Biotech). Primer synthesis and DNA sequencing were performed by Integrated DNA Technologies and Genewiz, respectively. PCR amplifications were performed on a Bio-Rad T100TM Thermal Cycler. Centrifugal filtration units were purchased from Millipore, and MINI dialysis units purchased from Pierce. Size exclusion chromatography was performed on an AKTA FPLC system from GE Healthcare equipped with a P-920 pump and UPC-900 monitor. Sephacryl S-200 columns were obtained from GE Healthcare. All the western blots were performed using the primary antibodies annotated in **Supplementary Table 3** and fluorophore-labeled secondary antibodies annotated in **Supplementary Table 4** following protocols recommended by the manufacture. Blots were imaged on an Odyssey CLx Imaging System (Li-Cor). Amino acid derivatives and coupling reagents were purchased from AGTC Bioproducts. Dimethylformamide (DMF), dichloromethane (DCM) and triisopropylsilane (TIS) were purchased from Fisher Scientific and used without further purification. Hydroxybenzotriazole (HOBt) and O-(benzotriazol-1-yl)-N,N,N′,N′-tetramethyluronium hexafluorophosphate (HBTU) were purchased from Fisher Scientific. Trifluoroacetic acid (TFA) was purchased from Fisher Scientific. N,Ndiisopropylethylamine (DIPEA) was purchased from Fisher Scientific. Analytical reversed-phase HPLC (RP-HPLC) was performed on an Agilent 1200 series instrument with an Agilent C18 column (5 μm, 4 × 150 mm), employing 0.1% TFA in water (HPLC solvent A), and 90% acetonitrile, 0.1% TFA in water (HPLC solvent B) as the mobile phases. Analytical gradients were 0-70% HPLC buffer B over 45 minutes at a flow rate of 0.5 mL/minute, unless stated otherwise. Preparative scale purifications were conducted on an Agilent LC system. An Agilent C18 preparative column (15-20 μm, 20 × 250 mm) or a semi-preparative column (12 μm, 10 mm × 250 mm) was employed at a flow rate of 20 mL/min or 4 mL/min, respectively. HPLC Electrospray ionization MS (HPLC-ESI-MS) analysis was performed on an Agilent 6120 Quadrupole LC/MS spectrometer (Agilent Technologies). All immunoblotting experiments in this research were performed at least 3X.

### Recombinant histone expression and purification

Recombinant human histones H2A, H2B, H3.2 and H4 were expressed in *E. coli* BL21 (DE3) or *E. coli* C41 (DE3), extracted by guanidine hydrochloride and purified by flash reverse chromatography, as previously described (1). The purified histones were analyzed by RP-LC-ESI-MS (1).

### Preparation of histone octamers and ‘601’ DNA

Octamers were prepared as previously described (1). Briefly, recombinant histones were dissolved in unfolding buffer (20 mM Tris-HCl, 6M GdmCl, 0.5mM DTT, pH 7.5), and combined with the following stoichiometry: 1.1 eq. H2A, 1.1 eq. H2B, 1 eq. H3.2, 1 eq. H4. The combined histone solution was adjusted to 1 mg/mL concentration and transferred to a dialysis cassette with a 7000 Da molecular cutoff. Octamers were assembled by dialysis at 4 °C against 3 × 1 L of octamer refolding buffer (10 mM Tris-HCl, 2 M NaCl, 0.5mM EDTA, 1 mM DTT, pH 7.5) and subsequently purified by size exclusion chromatography on a Superdex S200 10/300 column. Fractions containing octamers were combined, concentrated, diluted with glycerol to a final 50% v/v and stored at -20 °C. The 147-bp 601 DNA fragment was prepared by digestion from a plasmid containing 30 copies of the desired sequence (flanked by blunt EcoRV sites on either site) and purified by PEG-6000 precipitation as described before (1).

### Mononucleosome assembly

The mononucleosome assembly was performed according to the previously described salt dilution method with slight modification (1). Briefly, the purified wild-type octamers were mixed together with 601 DNA (1:1 ratio) in a 2 M salt solution (10 mM Tris pH 7.5, 2 M NaCl, 1 mM EDTA, 1 mM DTT). After incubation at 37 °C for 15 min, the mixture was gradually diluted (9 × 15 min) at 30 °C by dilution buffer (10 mM Tris pH 7.5, 10 mM NaCl, 1 mM EDTA, 1 mM DTT). The assembled mononucleosomes were concentrated and characterized by native gel electrophoresis (5% acrylamide gel, 0.5 × TBE, 120 V, 40 minutes) using ethidium bromide (EtBr) staining.

### Expression of recombinant wild-type TGM2 and C277A mutant

The pHis-hTGM2 plasmid was gifted from Byung Il Lee (Addgene plasmid #100719). The His_6_-tagged TGM2 C277A mutation was cloned by site-directed mutagenesis using pHis-hTGM2 as the template and the following primer sequences: 5′-GTCAAGTATGGCCAG<u>GCC</u>TGGGTCTTCGCCGCC-3′ and 5′-GGCGGCGAAGACCCA<u>GGC</u>CTGGCCATACTTGAC-3′. The gene sequences were confirmed by using the sequencing primer: 5′-GATGACCAGGGTGTGCTGCTG-3′. The His_8_-tagged wild-type and mutant TGM2 proteins were expressed in *E. coli* Rosetta (DE3) cells with an overnight IPTG induction at 16 °C. The bacterial pellet was lysed by sonication and the lysate was cleared by centrifugation at 12,000 r.p.m. for 30 minutes. The lysate was loaded on HisTrap HP Column (GE Healthcare) and eluted on the AKTA FPLC, followed by desalting using Zeba Spin Desalting Columns (7 K MWCO, 10 mL) according to the manufacturer’s protocol. Purified recombinant proteins were analyzed by SDS-PAGE and concentrated using stirred ultrafiltration cells (Millipore) according to the manufacturer’s protocol. The concentration of each protein was determined using 280 nm wavelength on a NanoDrop 2000c (Thermo Scientific).

### Peptide synthesis

Standard Fmoc-based Solid Phase Peptide Synthesis (FmocSPPS) was used for the synthesis of peptides in this study. Generally, the peptides were synthesized on ChemMatrix resins with Rink Amide to generate C-terminal amides. Peptides were synthesized using manual addition of the reagents (using a stream of dry N_2_ to agitate the reaction mixture). For amino acid coupling, 5 eq. Fmoc protected amino acid were pre-activated with 4.9 eq. HBTU, 5 eq. HOBt and 10 eq. DIPEA in DMF and then reacted with the N-terminally deprotected peptidyl resin. Fmoc deprotection was performed in an excess of 20% (v/v) piperidine in DMF, and the deprotected peptidyl resin was washed thoroughly with DMF to remove trace piperidine. Cleavage from the resin and side-chain deprotection were performed with 95 % TFA, 2.5% TIS and 2.5% H_2_O at room temperature for 1.5 hours. The peptides were then precipitated with cold diethyl ether, isolated by centrifugation and dissolved in water with 0.1 % TFA followed by RP-HPLC and ESI-MS analyses. Preparative RP-HPLC was used to purify the peptides of interest.

For the synthesis of site-specific monoaminylated H3 peptides in this study, Fmoc-Glu(OAII)-OH was incorporated at position 5 for the orthogonal deprotection and further monoaminylation. Briefly, the peptides were deprotected by Pd(PPh_3_)_4_ and PhSiH_3_ on resins (2) and then conjugated with monoamine donors [*i.e.*, histamine hydrochloride, serotonin hydrochloride (3), and acetonide-protected dopamine (4)] via PyAOP and DIEA catalysis (5).

For the synthesis of modified H3 peptide antigens in this study (as shown in **Supplementary Figure 6A**), Fmoc-Glu(OAII)-OH was incorporated at position 5 and either Fmoc-Lys(Boc)-OH or Fmoc-Lys(Me3)-OH was incorporated at position 4 on 2-Cl trityl resin via iterative FmocSPPS (i). The deallylation was conducted using Pd(PPh_3_)_4_ and PhSiH_3_ (ii), followed by the coupling of Trt-protected histamine (iii) and then acidolytic cleavage from the resin as well as global deprotection (iv). Note that both the validated H3Q5his and H3K4me3Q5his antibodies were licensed to Millipore for sale (see below for catalog numbers).

### *In vitro* biochemical assays

The TGM2 (de)monoaminylation assays were generally performed in the buffer (pH 7.5) containing 50 mM Tris-HCl, 5 mM CaCl_2_, and 2 mM DTT (freshly added). For H3 peptide (de)monoaminylation and monoamine-replacement, 2 mM peptides were treated with 100 μM TGM2 (or cell lysates) at 37 °C in the presence (or absence) of the corresponding monoamaines (4 mM) for 2 hours and then analyzed by LC-MS. For nucleosome core particle (NCP) (de)monoaminylation and monoamine-replacement, 1 μM NCPs were treated with 0.1 μM TGM2 at 37 °C in the presence (or absence) of the corresponding monoamaines (0.5 μM) for 2 hours. The (de)monoaminylated NCPs were analyzed by sodium dodecyl sulfate polyacrylamide gel electrophoresis (SDS-PAGE) followed by western blot analysis. H3 was used as the loading control in SDS-PAGE and western blot analyses. The buffer exchange for monoamine-replacement assays was performed using 0.5 mL Centrifugal Filter (3K, Millipore) with a 120-fold v/v for the removal of excess monoamine from the old reaction buffer systems.

### Expression of TGM2 in HEK 293T cells

The pShooter pCMV-nuc-myc vector expressing NLS-tagged wild type human TGM2 was utilized in our previous research (3). The catalytically dead mutant TGM2-C277A plasmid was constructed by site-directed mutagenesis using 5′-GTCAAGTATGGCCAG<u>GCC</u>TGGGTCTTCGCCGCC-3′ and 5′-GGCGGCGAAGACCCA<u>GGC</u>CTGGCCATACTTGAC-3′ as primers. The gene sequences were confirmed by using the sequencing primer: 5′-GATGACCAGGGTGTGCTGCTG-3′. Wild type TGM2 and the TGM2-C277A mutant were overexpressed in HEK 293T cells using Lipofectamine 2000 Transfection Reagent (Thermo Fisher Scientific) according to the manufacturer’s protocol. HEK 293T cells (ATCC) were cultured at 37 °C with 5% CO_2_ in DMEM medium supplemented with 10% fetal bovine serum (FBS) (Sigma-Aldrich), 2 mM L-glutamine and 500 units mL^−1^ penicillin and streptomycin. The cells were stimulated with 2 μM calcium ionophore (Sigma-Aldrich, A23187) for 6 h at 37 °C before lysis in DPBS buffer (Gibco), and then the expression of TGM2 was detected by western blot analyses with anti-TGM2 antibody (CST, #3557).

### Immunoprecipitation and pull-down

To capture the TGM2-H3 thioester complex, 50 μM of free H3 proteins were treated with 50 μM of wild type TGM2 or TGM2-C277A mutant in the buffer (pH 7.0) containing 50 mM Tris-HCl, 5 mM CaCl_2_, and 2 mM DTT (freshly added) at 37 °C for 1 hour. His_8_-tagged TGM2 was first pulled down by BSA-blocked Ni^2+^-NTA agarose beads (Thermo Fisher Scientific). Next, the beads were washed 3 times with Tris-HCl buffer (pH 7.0), boiled, separated on SDS-PAGE and analyzed by western blot with anti-TGM2 and anti-H3 antibodies to detect the enrichment of H3.

### Salt extraction of histones from cells

The extraction of histones from cells was performed according to the previously described high salt extraction method (1). Briefly, the cell lysis solution was prepared using extraction buffer (10 mM HEPES pH 7.9, 10 mM KCl, 1.5 mM MgCl_2_, 0.34 M sucrose, 10% glycerol, 0.2% NP40, protease and phosphatase inhibitors to 1 × from stock). After spinning down, the pellet was extracted using a no-salt buffer (3 mM EDTA, 0.2 mM EGTA). After discarding the supernatant, the final pellet was extracted by using high-salt buffer (50 mM Tris pH 8.0, 2.5 M NaCl, 0.05% NP40) in 4 °C cold room for 1 hour. After spinning down, the supernatant containing extracted histones was collected for further analyses.

### Cell fractionation

Cytosolic and nuclear fractions were prepared using NEPER Nuclear and Cytoplasmic Extraction Reagents (Thermo Scientific) according to the manufacturer’s protocol. Histones were extracted from the pellet using high salt extraction protocol, as described above (1). Purity of fractionation was evaluated using the following antibodies: anti-Actin (cytosol), anti-MEK ½ (nucleoplasm) and anti-H3 (chromatin).

### Pulse-chase experiments and inhibitor treatment

HEK 293T and HeLa cells were cultured at 37 °C with 5% CO_2_ in DMEM medium supplemented with 10% fetal bovine serum (FBS) (Sigma-Aldrich), 2 mM L-glutamine and 500 units mL^−1^ penicillin and streptomycin. Wild type TGM2 or TGM2-C277A mutant was overexpressed in 293T cells. The cultured cells were incubated with 500 μM monoamines (5-PT or histamine) for 6 h before the medium was changed to monoamine-free DMEM. Cells were cultured for an additional 6 hours, 12 hours, or 18 hours, after which they were washed by DPBS and harvested, and the cytosolic and histone fractions were prepared as described above (1, 6). Samples were separated on a single SDS-PAGE, transferred to a PVDF membrane and blotted with the indicated antibodies (or Cy5 dye) for further analyses. The cells were stimulated with 2 μM calcium ionophore (Sigma-Aldrich, A23187) during incubation with monoamine donors as described above.

For the inhibitor treatment assays, 100 µM of TGM2 inhibitors (ERW1041E or ZDON) were added to cell media 2 hours prior to adding the corresponding monoamine donors. Cells were incubated for additional 6 hours after which they were harvested, and histones were extracted and analyzed as described above. Samples were separated on a single SDS-PAGE, transferred to a PVDF membrane and blotted with the indicated antibodies. For the inhibitor-treated *in vitro* biochemical assays, where 1 μM NCPs were used as substrate and 0.1 μM TGM2 was used as catalyst, 1 μM ERW1041E was added to the reaction systems for inhibiting the activity of TGM2.

### Visualization of H3 serotonylation by CuAAC

HEK 293T and HeLa cells were treated with 500 μM 5-PT and stimulated with 2 μM calcium ionophore (Sigma-Aldrich, A23187) during incubation, as described above. The cells were washed by DPBS, harvested and then the histone factions were extracted via high salt extraction. Extracted histones were desalinated, lyophilized and then resuspended in DPBS buffer containing 0.4 % SDS. 50 μL of freshly dissolved histones was added to a premixed solution containing 3 μL of 10 mM Cy5-azide (Sigma-Aldrich, 777323), 10 μL of a 3:7 mixture of 50 mM CuSO_4_ and 100 mM THPTA, and then vortexed. Thereafter, 5 μL of 100 mM freshly made TCEP was added to initiate the click reaction followed by incubation (1-2 hours) at 30 °C. Then, 10 μL of 0.5 M EDTA was added to quench the reactions. Excess reagents were removed by MeOH/CHCl_3_ protein precipitation or concentration-dilution using a 0.5 mL Centrifugal Filter (3K, Millipore). Pellets were washed by 500 μL MeOH/H2O (9:1) prior to a second centrifugation (7). The air-dried protein samples were then analyzed by SDS-PAGE followed by in-gel imaging using Odyssey CLx Imaging System (wavelength 680 nm).

### Identification of H3 histaminylation from cells

HEK 293T cells transfected with wild type TGM2 or TGM2-C277A mutant plasmids were treated with 500 μM histamine as described above. Histones were then extracted from harvested cell pellets, as described before (8), for further mass spectrometry analysis. Briefly, histones were extracted with chilled 0.2 M sulfuric acid (5:1, sulfuric acid : pellet) and incubated with constant rotation for 4 hours at 4 °C, followed by precipitation with 33% trichloroacetic acid overnight at 4°C. Then, the supernatant was removed and the tubes were rinsed with ice-cold acetone containing 0.1% hydrochloric acid, centrifuged and rinsed again using 100% ice-cold acetone. After centrifugation, the supernatant was discarded and the pellet was dried using lyophilizer. The pellet was dissolved in 50 mM ammonium bicarbonate (pH 8.0), and histones were subjected to derivatization using 5 µL of propionic anhydride and 14 µL of ammonium hydroxide (Sigma-Aldrich) to balance the pH at 8.0. The mixture was incubated for 15 min and the procedure was repeated. Histones were then digested with 1 µg of sequencing grade trypsin (Promega) diluted in 50mM ammonium bicarbonate (1:20, enzyme : sample) overnight at room temperature. Derivatization reaction was repeated to derivatize peptide N-termini. The samples were dried by lyophilizer. Prior to mass spectrometry analysis, samples were desalted using a 96-well plate filter (Orochem) packed with 1 mg of Oasis HLB C-18 resin (Waters). Briefly, the samples were resuspended in 100 µl of 0.1% TFA and loaded onto the HLB resin, which was previously equilibrated using 100 µl of the same buffer. After washing with 100 µl of 0.1% TFA, the samples were eluted with a buffer containing 70 µl of 60% acetonitrile and 0.1% TFA and then dried by lyophilizer.

Samples were analyzed using nano liquid chromatography coupled online with tandem mass spectrometry (nLC-MS/MS). Briefly, samples were resuspended in 10 µl of 0.1% trifluoroacetic acid (TFA) and loaded onto a Dionex RSLC Ultimate 300 (Thermo Scientific), coupled online with an Orbitrap Fusion Lumos (Thermo Scientific). Chromatographic separation was performed with a two-column system, consisting of a C-18 trap cartridge (300 µm ID, 5 mm length) and a picofrit analytical column (75 µm ID, 25 cm length) packed in-house with reversed-phase Repro-Sil Pur C18-AQ 3 µm resin. Histone peptides were separated using a 30 min gradient from 4-30% buffer B (buffer A: 0.1% formic acid; buffer B: 80% acetonitrile + 0.1% formic acid) at a flow rate of 300 ml/min. The mass spectrometer was set to acquire spectra in a data-independent acquisition (DIA) mode. The full MS scan was set to 300-1100 m/z in the orbitrap with a resolution of 120,000 (at 200 m/z) and an AGC target of 5x10e5. MS/MS was performed in the orbitrap with sequential isolation windows of 50 m/z with an AGC target of 2x10e5 and an HCD collision energy of 30.

Targeted MS/MS was performed for the endogenous Q5his peptide (m/z 455.7587). Data were manually inspected and the peak intensity was obtained by calculating the area of the extracted ion chromatogram. Histone peptides raw files were imported into EpiProfile 2.0 software (9) to quantify acetylated, methylated and phosphorylated peptides. From the extracted ion chromatogram, the area under the curve was obtained and used to estimate the abundance of each peptide. In order to achieve the relative abundance of post-translational modifications, the sum of all different modified forms of a histone peptide was considered as 100%, and the area of the particular peptide was divided by the total area for that histone peptide in all of its modified forms. The relative ratio of two isobaric forms was estimated by averaging the ratio for each fragment ion with different mass between two species. The resulting peptide list generated by EpiProfile was exported to Microsoft Excel and further processed for a detailed analysis.

### Animals

Mice (C57BL/6J) were purchased from The Jackson Laboratory. Animals were group housed (2-5 per cage) on a 12-hour light/dark cycle (lights on from 7:00 A.M. to 7:00 P.M.) at constant temperature (23°C) with *ad libitum* access to food and water. All animal protocols were approved by the IACUC at both the Icahn School of Medicine at Mount Sinai (ISMMS).

### Immunoblotting - Brain

Brain tissues were extracted from euthanized mice and immediately frozen whole. Brains were later sectioned using razor blades and a brain block to 1-mm thickness, with tissue punches (1-2 mm) collected for corresponding brain regions. To purify nuclear fractions, punches were homogenized in Buffer A containing 10 mM HEPES (pH 7.9), 10 mM KCl, 1.5 mM MgCl_2_, 0.34 M sucrose, 10% glycerol, 1 mM EDTA, and 1X protease inhibitor cocktail. Following homogenization, 0.1% TritonX-100 was added to each homogenate, incubated and rotated at 4 °C for 30 minutes and then centrifuged for 5 min at 1300g at 4° C. Supernatants containing cytosolic fractions were discarded, and nuclear pellets were resuspended in Buffer A to remove any remaining cytosolic contamination, followed by centrifugation for 5 minutes at 1300g at 4 °C. After centrifugation, supernatants were discarded and pellets were resuspended and sonicated in sample buffer containing 0.3 M sucrose, 5 mM HEPES, 1% SDS and 1X protease inhibitor cocktail. Protein concentrations were measured using the DC protein assay kit (BioRad), and 1-20 µg of protein was loaded onto 4-12% NuPage BisTris gels (Invitrogen) for electrophoresis. Proteins were then transferred to PVDF membranes and blocked for 30 minutes in 5% milk in PBS + 0.1% Tween 20 (PBST), followed by incubation with primary antibodies overnight at 4 °C. For competition assays, antibodies were pre-incubated with indicated peptides at a 5:1 ratio for 1 hour at RT before being incubated with the membrane. The following antibodies were used: rabbit anti-H3Q5his (1:200, Millipore ABE2578), rabbit anti-H3K4me3Q5his (1:500, Millipore ABE2605), rabbit anti-H3K4me3 (1:1000, Abcam ab8580), rabbit anti-H3K4me2 (1:1000, Abcam ab7766), rabbit anti-H3K4me1 (1:1000, Abcam ab8895), rabbit anti-Tgm2 (1:500, Cell Signaling 3557S), and rabbit anti-H3 (1:50000, Abcam ab1791). The next day, membranes were washed 3X in PBST (10 minutes) and incubated for 1 hour with horseradish peroxidase conjugated anti-rabbit secondary antibody (BioRad 170-6515; 1:10000; 1:50000 for anti-H3 antibody) in 5% milk/PBST at RT. After three final washes with PBST, bands were detected using enhanced chemiluminescence (ECL; Millipore). Densitometry was used to quantify protein bands using Image J Software (NIH), and proteins were normalized to total H3 or H4. For peptide dot blots, peptides (unmodified *vs.* H3Q5his *vs.* H3Q5ser *vs.* H3Q5dop) were dotted as progressive protein concentrations (0.25, 0.5, 1 µg) on a nitrocellulose membrane. Membranes were left to dry at RT for 1 hour and then blocked in 5% milk/PBST for 1 hour. Membranes were treated similar to that described above.

### Enzymatic assays for antibody validations

For assessments of TGM2-mediated transamidation of histamine to histone H3, 0.25 µg guinea pig Tgm2 (Zedira, T006), 5 mM histamine (Sigma) and 10 µg of recombinant H3.2 were combined with enzymatic buffer containing 250 mM tris-acetate (pH 7.5), 8.75mM CaCl2 and 1X protease inhibitor cocktail, followed by incubation for 3 hours at RT. Following incubation, enzymatic reactions were boiled with laemmli buffer and then run on 4-12% NuPage BisTris gels (Invitrogen) and blotted as described previously. Enzymatic assays were also performed using serotonin and dopamine to confirm the specificity of our in-house anti-H3Q5his antibody. Enzymatic assays were also performed using reconstituted unmodified vs. K4me3 mononucleosomes from Active Motif.

### LC-MS/MS validation of histaminylated H3 (1–15)

Samples were analyzed by LC-MS/MS (Dionex 3000 coupled to Q-Exactive mass spectrometer, Thermo Fisher). Peptides were separated by C-18 reversed phase chromatography (inner diameter of 75 µm, particle size of 3µm, Nikkyo Technologies, Japan) using a gradient increasing from 1% B to 25% B in 16 minutes (A: 0.1% formic acid, B: 80% acetonitrile in 0.1% formic acid). The mass spectrometer was operated in parallel reaction monitoring (PRM) mode https://www.ncbi.nlm.nih.gov/pmc/articles/PMC6557285/ - R35 (MS and MS/MS resolution of 70,000 and 35,000 respectively) (10) with an AGC target, 5e5, max. inject time of 60ms and an isolation m/z window of 1.3). MS was acquired from m/z 300– 1650 while m/z 100 was set as lowest mass for MS/MS. Charge states 2+ to 5+ of the modified peptide: (ARTKQ(histamine)TARKSTGGKA-NH_2_), were targeted. An energy of 25 NCE was used for the peptide in charge states 3+ and 4+ while NCE of 35 was used for the doubly charge version. **Supplementary Figure 5D** represents a high resolution, high accuracy tandem mass spectra (sample: +histamine/+TGM2) of the doubly charged peptide: ARTKQTARKSTGGKA-NH2 modified by histamine at Gln 5 (m/z 551.9930 [1.5 ppm]). The full amino acid sequence was accounted for. Selected fragment ions, including y10/b5 and y11/ b4 that identify the histamine modified glutamine, are annotated.

### AAV constructs and viral transduction

Adeno-associated virus H3.3 constructs [empty vs. wildtype (WT) vs. (Q5A)-Flag-HA] were generated and validated, as previously described (11). All three vectors contain an IRES driven GFP fluorescent tag to allow visualization of the injection site during tissue dissection. Animals were anaesthetized with isoflurane (1-3%) and positioned in a stereotaxic frame (Kopf instruments) and 0.5 μl of viral construct was infused bilaterally into TMN using the following coordinates; anterior-posterior (AP) -2.3, medial-lateral (ML) +1.0, dorsal-ventral (DV) -5.5. Following surgery, mice received meloxicam (1 mg/kg) s.c. and topical antibiotic treatments for 3 days. All tissue collections or behavioral testing commenced 21 days after surgery to allow for both maximal expression of the viral constructs.

### Immunohistochemistry

Mice were anaesthetized with ketamine/xylazine (100/12 mg/kg) i.p., and then perfused transcardially with cold phosphate buffered saline (PBS 1X) followed by 4% paraformaldehyde (PFA) in 1X PBS. Next, brains were post-fixed in 4% PFA overnight at 4° C and then transferred into 30% sucrose/PBS 1X for two days. Brains were then cut into serial 40 μm coronal slices. Free floating TMN slices were washed 3X in tris buffered saline (TBS 1X), incubated for 30 minutes in 0.2% Triton X/TBS 1X–to permablilize tissue–and then incubated for 1 hour at RT in blocking buffer (0.3% Triton X, 3% donkey serum, 1X TBS). Brain slices were then incubated overnight at RT with mouse anti-GFP (1:200; Abcam ab65856) and HA-488 (1:200; Life Technologies, Alexa Fluor SC-805). The next day, brain slices were washed 3X in 1X TBS and then incubated for 2 hours at RT with a fluorescent-tagged Alexa Fluor 568 anti-mouse secondary antibody (1:500; Life Technologies A11004). Brain sections were then washed 3X in 1X TBS, incubated with DAPI (1:10000, Thermo Scientific 62248) for 5 minutes at RT, mounted on Superfrost Plus slides (Fischer Scientific) and then coverslipped with Prolong Gold (Invitrogen). Immunofluorescence was visualized using a confocal microscope (Zeiss LSM 780).

### RNA-sequencing

Brain tissue was collected every 4 hours across the 24 hour zeitgeber, 21 days post viral-infusion, or following behavioral assays. All brains were immediately frozen following collection. For zeitgeber brains, tissues were sectioned at 1 mm thick and TMN was collected by tissue punch (1-2 mm). For virus-infused brains, tissue was sectioned at 150 μm on a cryostat, and GFP was illuminated using a NIGHTSEA BlueStar flashlight to microdissect virally infected tissues. TMN tissue punches were homogenized in Trizol (Thermo Fisher), and RNA was isolated on RNeasy Microcolumns (Qiagen) following manufacturer’s instructions. RNA concentrations were assessed using a Qubit spectrophotometer and quality was confirmed using a bioanalyzer. Following RNA purification, RNA-seq libraries were prepared according to Illumina protocols and sequenced on an Illumina HiSeq 2500 (or equivalent) sequencer. Raw sequencing reads from mice were mapped to mm10 using HISAT2 (12). Counts of reads mapping to genes were obtained using htseq-counts software (13) against Ensembl v90 annotation. fpkm function in DESeq2 (14) was utilized to obtain the FPKM values of each gene. The R program JTK_CYCLE (15) was utilized to identify cycling genes from the dataset (ADJ.P<0.05). pheatmap package in R (16) was used to generate a heatmap depicting the cyclic genes. Cyclic genes were further assessed using ChEA (17) in Enrichr (https://maayanlab.cloud/Enrichr/), which infers transcription factor regulation from integration of previous genome-wide chromatin immunoprecipitation (ChIP) analyses. For viral transduction RNA-seq studies (collected at ZT 20), sources of technical variation between samples were calculated for the combined control viral groups (GFP + H3.3 WT) *vs.* Q5A using the RUVg (k=5) model of RUVseq (Version 4.1.1). These factors of variation were incorporated into the design formula for DESEQ2 (14) (Version 2.11.40.6) in order to perform pairwise differential expression analyses between Q5A *vs.* control viral groups. Differentially expressed (DE) genes were defined at FDR < .05 (n=1470). Odds Ratio analyses were carried out on this DE gene list against our previously identified list of circadian rhythmic genes (n=2338), as well as against a set of verified CLOCK transcription factor target genes from the CHEA database (n=387) (17) using the GeneOverlap R package (Version 1.26.0) (18).

### Locomotor activity

In order to monitor locomotor activity across sleep:wake cycles, mice were individually placed in clean, transparent home cages with minimal bedding and access to food and water *ad libitum*. Home cages were placed in larger activity chambers with infrared beams to detect movement across 24 hours (clear plexiglass 40 × 40 × 30 cm, Omnitech Electronics Inc., Columbus, OH), starting at 19:00 (lights off). Activity was monitored via beam breaks, which was collected by Fusion Software (v5.0) (Omnitech Electronics) software and calculated into 4-hour time bins. Animals were monitored 12 hours after switching them from their normal light:dark cycle (lights on at 7:00, lights off at 19:00) to dark:dark for 48 hours in order to assess their locomotor behavior.

### Sleep manipulation

Mice were individually habituated to locomotor activity monitoring cages and chambers (as described previously in ‘Locomotor Activity’ methods) for 24 hours, and then received either an injection of vehicle or zolpidem (10 mg/kg, i.p.) at 19:00 (beginning of active phase, zeitgeber 12). Animals were placed back in the locomotor activity monitoring cages immediately and activity was measured for 8 hours. Animals were immediately euthanized and brains were collected and frozen for subsequent immunoblotting experiments.

### Recombinant protein cloning and purification of WDR5-WT, WDR5-mut, MLL1 complex

Full-length WDR5 and truncated WDR5 residues 22 to 334 (WDR5^22–334^) were cloned into pET28b vector for protein purification. Full-length WDR5-K259A and WDR5-K259E mutants were generated by Site-Directed Mutagenesis Kit (Agilent). All proteins were expressed in the *E. coli* BL21 DE3 (Novagen) and induced overnight by 0.2 mM isopropyl β-D-thiogalactoside at 16 °C in the LB medium. The collected cells were suspended in 500 mM NaCl, 20 mM Tris, pH 7.5. After cell lysis and centrifugation, the supernatant was applied to HisTrap column (GE Healthcare). After washing 5 column volumes with the suspension buffer, the protein was eluted with the buffer 100 mM NaCl, 20 mM Tris pH 7.5, 500 mM imidazole and cut with Thrombin enzyme overnight. Proteins were further purified by the HiTrap SP (GE Healthcare) cation-exchange column and a HiLoad 16/60 Superdex 75 (GE Healthcare) gel filtration column using AKTA Purifier 10 systems (GE Healthcare). All proteins were stored in 100 mM NaCl, 20 mM Tris, pH 7.5 at ∼ 10 mg/ml in an -80°C freezer. Human MLL1 constructs (3745–3969), as well as full-length human WDR5, RBBP5, DPY30 and ASH2L (95–628) proteins, were individually expressed in E. coli BL21 cells. All proteins were induced overnight with 0.2 mM isopropyl β-D-thiogalactoside at 16°C in LB medium. Cell pellets were suspended in lysis buffer (20 mM Tris-HCl, pH 8.0, 500 mM NaCl, 5% glycerol, 1 mM DTT). After cell lysis and centrifugation, supernatants were purified using HisTrap columns (GE Healthcare) or Glutathione Sepharose 4B beads (GE Healthcare), followed by enzyme digestion to remove tags. All proteins were further purified on HiTrap SP (GE Healthcare) cation-exchange columns or HiTrap Q (GE Healthcare) anion-exchange columns. MLL1, WDR5 and DPY30 were further purified with the HiLoad 10/300 Superdex 75, while ASH2L and RBBP5 were further purified with the HiLoad 10/300 Superdex 200. The buffer for gel-filtration chromatography contained 50 mM Tris-HCl, pH 7.5, 300 mM NaCl, 1 mM DTT and 10% glycerol. Purified proteins were concentrated to 10-20 mg/ml and stored at -80 °C. Note that for experiments presented in **Figure 5B-C**, MLL1 complex was purchased from Active Motif (catalog #31423).

### MALDI-TOF analysis of enzymatic activity

H3K4 methyltransferase assays were conducted by combining 1.2 μM of the MLL^3745-3969^-WDR5-RBBP5-ASH2L-DPY30 complex with 10 μM histone H3 peptide and 250 μM methyl-Sadenosyl-methionine in 50 mM Tris, pH 8.5, 50 mM KCl, 5 mM dithiothreitol, 5 mM MgCl_2_, and 5% glycerol at 15°C. Reactions were quenched by the addition of HPLC solvent A (H_2_O + 0.1% TFA) and were desalted using C18 ZipTip (Millipore) according to manufacturer’s protocol before being diluted 1:1 with α-cyano-4-hydroxycinnamic acid (CHCA) matrix in 50% ACN plus 20% acetone with 0.1% TFA and spotted on a MALDI-TOF plate for analysis. Samples were analyzed using a Bruker UltrafleXtreme MALDI TOF/TOF Mass Spectrometer and data were analyzed using Bruker Compass flexAnalysis Version 3.4.

### LC-MS/MS analysis of enzymatic activity

H3K4 methyltransferase assays were conducted by titrating concentrations of the MLL^3735-3973^-WDR5-RBBP5-ASH2L-DPY30 complex with 10 μM histone H3 peptide and 250 μM methyl-Sadenosyl-methionine in 50 mM Tris, pH 8.5, 50 mM KCl, 5 mM dithiothreitol, 5 mM MgCl_2_, and 5% glycerol at 15°C. Samples were diluted and quenched by adding 30 µL 0.1% trifluoro acetic acid (TFA) and subjected to a C18-based micro solid-phase extraction (19) to desalt and remove/minimize enzyme. Peptides were eluted using 40% acetonitrile in 0.1% TFA. Eluted samples were dried and re-dissolved in 20 µL 3% acetonitrile/0.1% formic acid. 5 µL were injected onto C18 column (Acclaim 120A C18 5um 2.1mm X 150mm) connected to an Orbitrap XL (Thermofisher). Peptides were separated using a gradient delivered at 200 µL/minute, increasing from 1% B/99% A to 45% B/55% A (B: acetonitrile/0.1%, formic acid, A: 0.1% formic acid) for 13.5 minutes. Masses of eluting analysts were measured at 60,000 resolution (high resolution/high mass accuracy). Data (MS1 signals) were processed using Skyline (20). The signals of the C-terminal amidated peptides were measured: ARTKQTARKSTGGKAPRKQLA, as well as the glutamine (Q) 5 histaminylated or serotonylated forms with and without mono-, di- and trimethylated Lysine (K) 4 residue. Peptides with charge states 2+ through 6+ were extracted.

### ITC

For ITC measurement, synthetic histone peptides and the full-length wild-type and mutant WDR5 proteins were extensively dialyzed against ITC buffer: 100 mM NaCl and 20 mM Tris, pH 7.5. The titration was performed using a MicroCal iTC200 system (GE Healthcare) at 25 °C. Each ITC titration consisted of 17 successive injections with 0.4 μl for the first and 2.4 μl for the rest. Peptides were titrated into proteins in all the experiments. The resultant ITC curves were processed using Origin 7.0 software (OriginLab) according to the ‘One Set of Sites’ fitting model.

### Histone tail peptide IPs against HeLa nuclear extracts and recombinant WDR5

Biotinylated unmodified *vs.* H3Q5ser *vs.* H3Q5his peptides (100 μg) were resuspended with 100 μL of prewashed immobilized Streptavidin beads (DynaBeads™ Streptavidin M-280) in 0.01% DPBS/Triton-X 100, with subsequent incubation (rotating) overnight at 4°C. For each IP, 40 μl of 50% peptide-bead slurry was added to either dialyzed HeLa cell nuclear extracts (10 mg/IP, with 2 mM MgCl_2_ added prior to the beads) or full length recombinant WDR5 (purified as described above, in 150 mM KCl, 50 mM Tris-HCl at pH 8.0, 1% NP40, 1 mM DTT), and each sample rotated for at 4°C for 4 hours. IPs were then centrifuged for 1 minute at 1000 rpm to pellet the beads. Beads were subsequently washed six times in binding buffer substituted with 300 mM KCl (and no MgCl_2_). For western blotting analyses, beads were then washed in cold DPBS and were boiled for 8 minutes in 30 μl of denaturing sample buffer.

### Crystallization and X-Ray structure determination

Truncated WDR^522–334^ was firstly incubated with H3K4me3Q5his peptide at a molar ratio 1:2 for 1 hour. Crystallization was performed by the sitting-drop vapor diffusion method under 18°C by mixing equal volumes (1-2μl) of protein and reservoir solution. The crystal was obtained at the condition 0.1 M sodium citrate tribasic dihydrate (pH 5.5), 22% polyethylene glycol (PEG) 3350, and 0.1% n-Octyl-β-D-glucoside at 18 °C. Crystals were briefly soaked in the cryo-protectant and were flash frozen in liquid nitrogen for data collection at 100K. Complex data sets were collected at beamline BL17U at the Shanghai Synchrotron Radiation Facility. All data were indexed, integrated and merged using the HKL2000 software package (21). The complex structures were solved by molecular replacement using MOLREP (22). All structures were refined using PHENIX (23), with iterative manual model building using COOT (24). Model geometry was analyzed with PROCHECK (25). The electron density of H3Q5 was visible while the histamine modification density was not clear. In the WDR5-H3Q5his structure and MLL3-RBBP5-ASH2L-H3 complex structure (PDB ID: 5F6K), the model of histamine was built based upon the orientation of H3Q5 residue and restricted within the Ramachandran plot. All structural figures were created using PYMOL (http://www.pymol.org/).

### Statistics and reproducibility

All *in vitro* and *in cellulo* western blotting and mass spectrometry analyses were repeated independently at least three times with similar results. For western blot comparisons in brain examining rhythmic patterns of expression for H3Q5his, H3K4me1/2/3, H3K4me3Q5his and TGM2 (all normalized to total H3), non-linear regression ‘comparison of fits’ analyses were performed between third order polynomial, cubic trends (alternative hypothesis; i.e., rhythmic), *vs.* first order polynomial, straight line trends (null hypothesis; non-rhythmic). For behavioral locomotor testing involving two viral treatments and multiple time points, Two-way ANOVAs were performed with subsequent Sidak’s post hoc analyses for multiple comparisons, as well as *a posteriori* Student’s t-tests (as indicated in the text). For molecular analyses in brain (e.g., western blotting) comparing only two groups, unpaired Student’s t-tests were performed. For molecular analyses in brain (e.g., western blotting) comparing more than two groups (e.g., Zolpidem blots), one-way ANOVAs were performed, followed by Dunnett’s post hoc analyses for multiple comparisons. For biochemical quantifications of MLL1 activity (LC-MS/MS) or binding interactions presented in Figure 5, two- or one-way ANOVAs were employed with Sidak’s or Tukey’s multiple comparison tests, respectively. All animals used were included as separate *n*s (samples were not pooled). Significance was determined at p<0.05. All data are represented as mean ± SEM. Statistical analyses were performed in GraphPad Prism 9.

## SUPPLEMENTARY TABLES

**Supplementary Table 1.** ITC statistics.

| Peptide | Protein | $K_a$<br>($10^4 \cdot M^{-1}$) | $\Delta H$<br>(kcal/mol) | $\Delta S$<br>(cal/mol/deg) | N |
| --- | --- | --- | --- | --- | --- |
| H3<sub>(1-15)</sub>Q5un | WDR5-WT | 2.34±0.28 | -4.85±0.60 | 3.71 | 0.85±0.09 |
|  | WDR5-K259A | 5.62±0.12 | -4.84±0.28 | 5.52 | 1.00 |
|  | WDR5-K259E | 5.25±0.90 | -4.22±0.33 | 7.45 | 1.27±0.07 |
| H3<sub>(1-15)</sub>Q5his | WDR5-WT | 0.82±0.07 | -6.96±0.33 | -5.43 | 1.00 |
|  | WDR5-K259A | 6.22±0.74 | -4.46±0.20 | 6.96 | 1.25±0.04 |
|  | WDR5-K259E | 13.7±0.81 | -3.59±0.05 | 11.5 | 1.07±0.01 |

**Supplementary Table 2.**
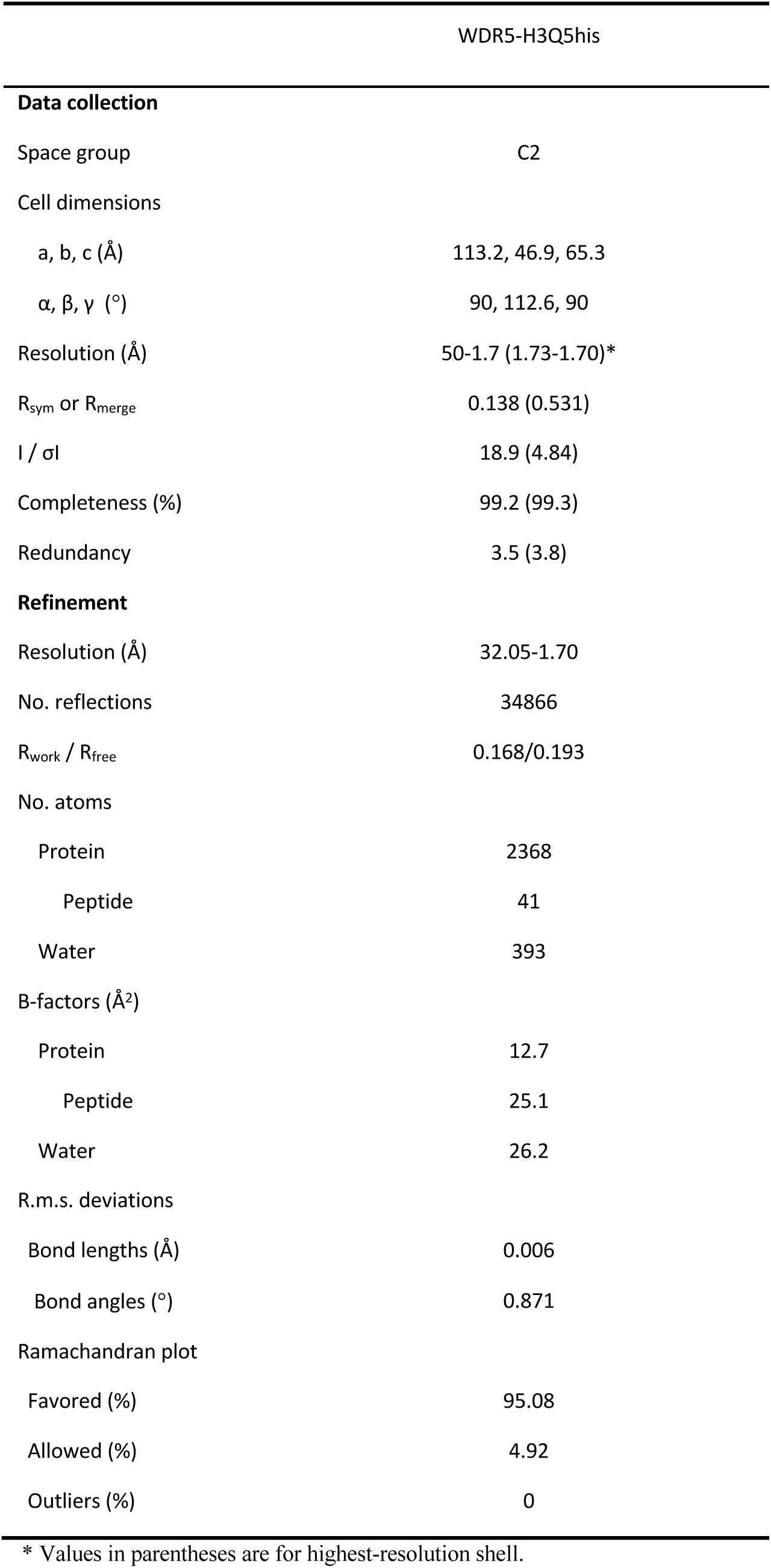
X-ray crystallography data collection and refinement statistics.

**Supplementary Table 3.** Primary antibodies used in this study.

| **Host** | **Epitope** | **WB** | **Vendor** |
| --- | --- | --- | --- |
| **Rabbit** | Anti-TGM2 | 1:500 (brain)<br>1: 1000 (in vitro and in cellulo) | CST<br>(3557S) |
| **Chicken** | Anti-H3 | 1: 1000 | Abcam<br>(ab134198) |
| **Mouse** | Anti-H3 | 1: 1000 | Abcam<br>(ab10799) |
| **Mouse** | Anti-Actin | 1: 1000 | CST<br>(3700S) |
| **Rabbit** | Anti-H3Q5ser | 1: 1000 | Millipore<br>(ABE1791) |
| **Rabbit** | Anti-H3Q5dop | 1: 1000 | Millipore<br>(ABE2588) |
| **Rabbit** | Anti-H3Q5his | 1: 200 (brain)<br>1:1000 (in vitro and in cellulo) | Millipore<br>(ABE2578) |
| **Rabbit** | Anti-H3K4me3Q5ser | 1:500 | Millipore<br>(ABE2605) |
| **Rabbit** | Anti-H3K4me3 | 1:1000 | Abcam<br>(ab8580) |
| **Rabbit** | Anti-H3K4me2 | 1:1000 | Abcam<br>(ab7766) |
| **Rabbit** | Anti-H3K4me1 | 1:1000 | Abcam<br>(ab8895) |
| **Rabbit** | Anti-H3 | 1:50000<br>(brain) | Abcam<br>(ab1791) |
| **Rabbit** | Anti-ASH2L | 1:1000 | Active Motif<br>(39099) |
| **Rabbit** | Anti-WDR5 | 1:1000 | CST<br>(13105) |
| **Rabbit** | Anti-RBBP5 | 1:1000 | Active Motif<br>(61405) |

**Supplementary Table 4.** Secondary antibodies used in this study.

| **Host** | **Epitope** | **Label** | **Dilution** | **Vendor** |
| --- | --- | --- | --- | --- |
| Donkey | Anti-Chicken | IRDye 800CW | 1: 15000 | Li-Cor |
| Goat | Anti-Mouse | IRDye 680RD | 1: 15000 | Li-Cor |
| Goat | Anti-Mouse | IRDye 800CW | 1: 15000 | Li-Cor |
| Goat | Anti-Rabbit | IRDye 800CW | 1: 15000 | Li-Cor |
| Goat | Anti-Rabbit | IRDye 680RD | 1: 15000 | Li-Cor |
| Goat | Anti-Rabbit | Horseradish Peroxidase | 1:10000<br>1:50000<br>(for anti-H3 antibody) | BioRad |
| Sheep | Anti-Mouse | Horseradish Peroxidase | 1:5000 | Cytiva |
| Donkey | Anti-Rabbit | Horseradish Peroxidase | 1:5000 | Cytiva |

## SUPPLEMENTARY FIGURES AND LEGENDS

**Supplementary Figure 1.**
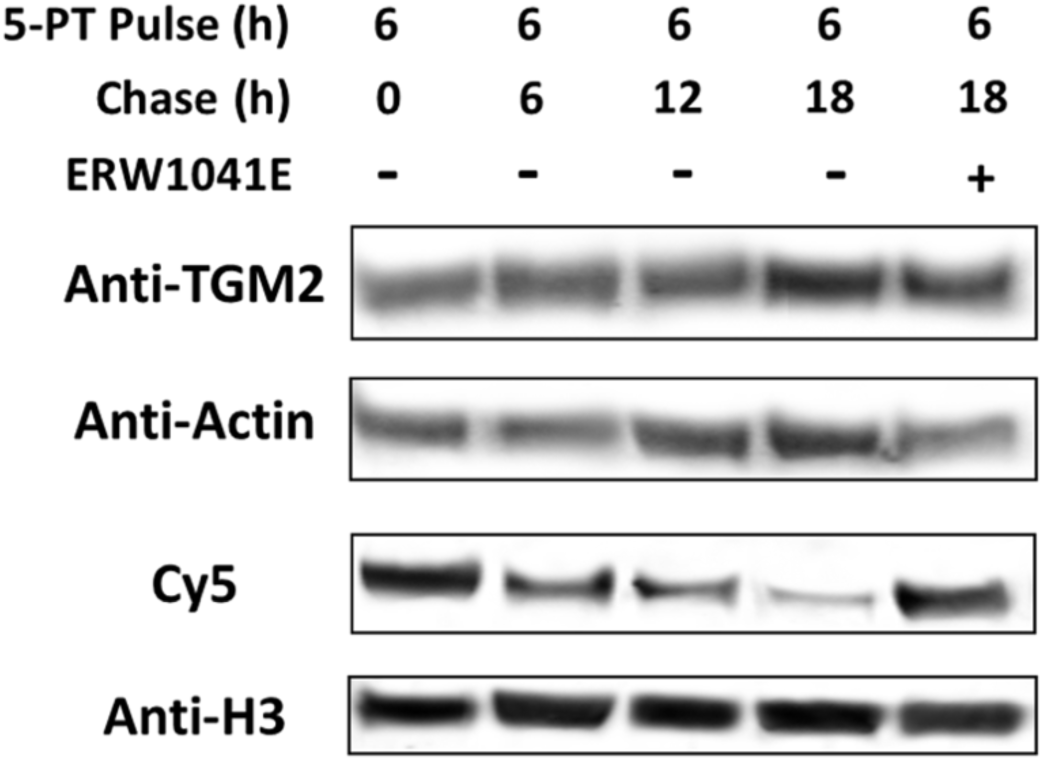
Time-course and pulse-chase analysis of histone H3 serotonylation. HeLa cells were treated with 500 μM 5-PT-treated and cultured for 6 h before the medium was changed to monoamine-free DMEM with or without 100 μM TGM2 inhibitor, ERW1041E. Cells were cultured for an additional 6 h, 12h or 18 h before being harvested. Histones were then extracted, labeled by Cy5 via CuAAC, separated by SDS-PAGE, transferred to a PVDF membrane and blotted with indicated antibodies. Experiment repeated >3X.

**Supplementary Figure 2.**
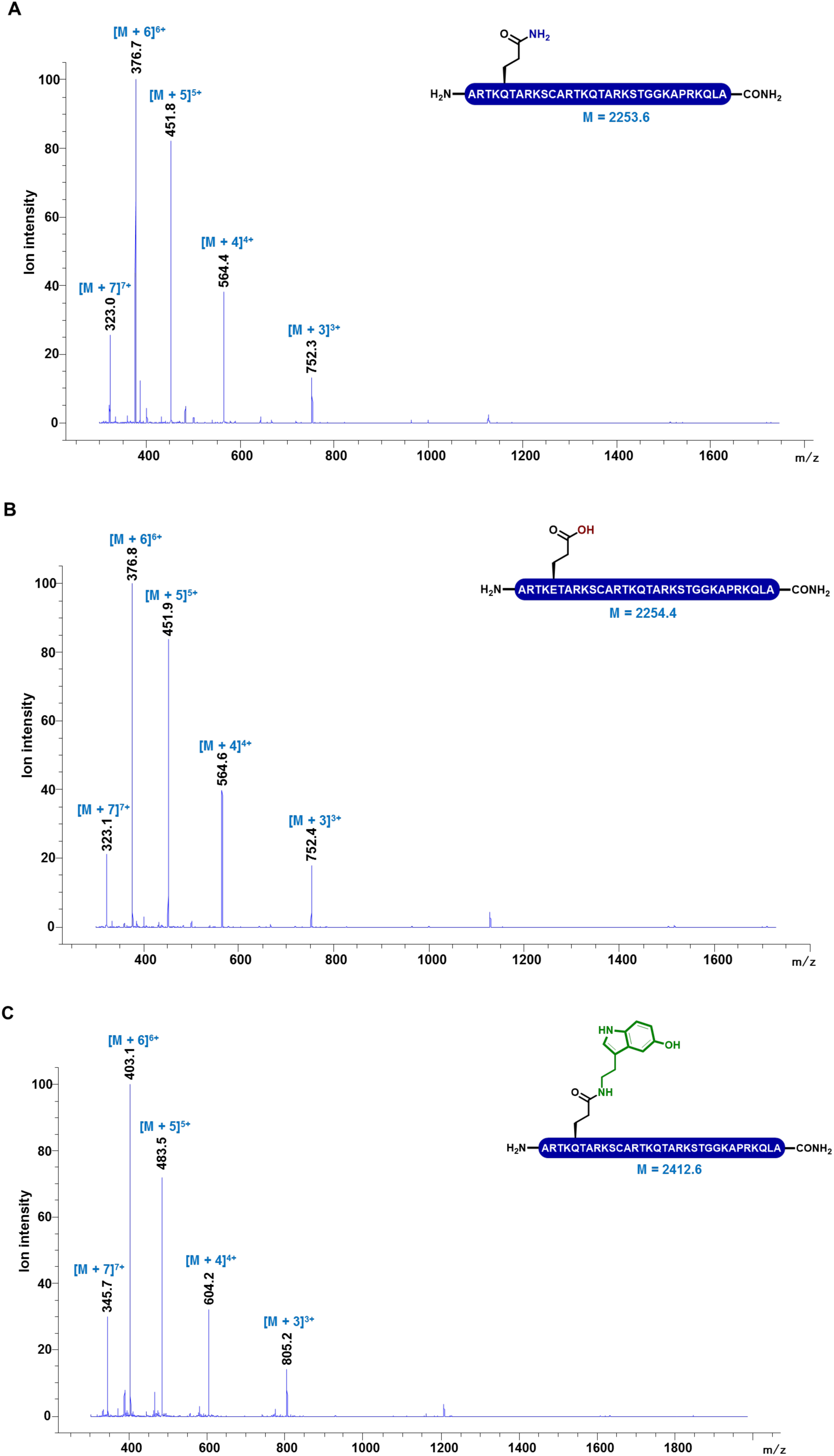
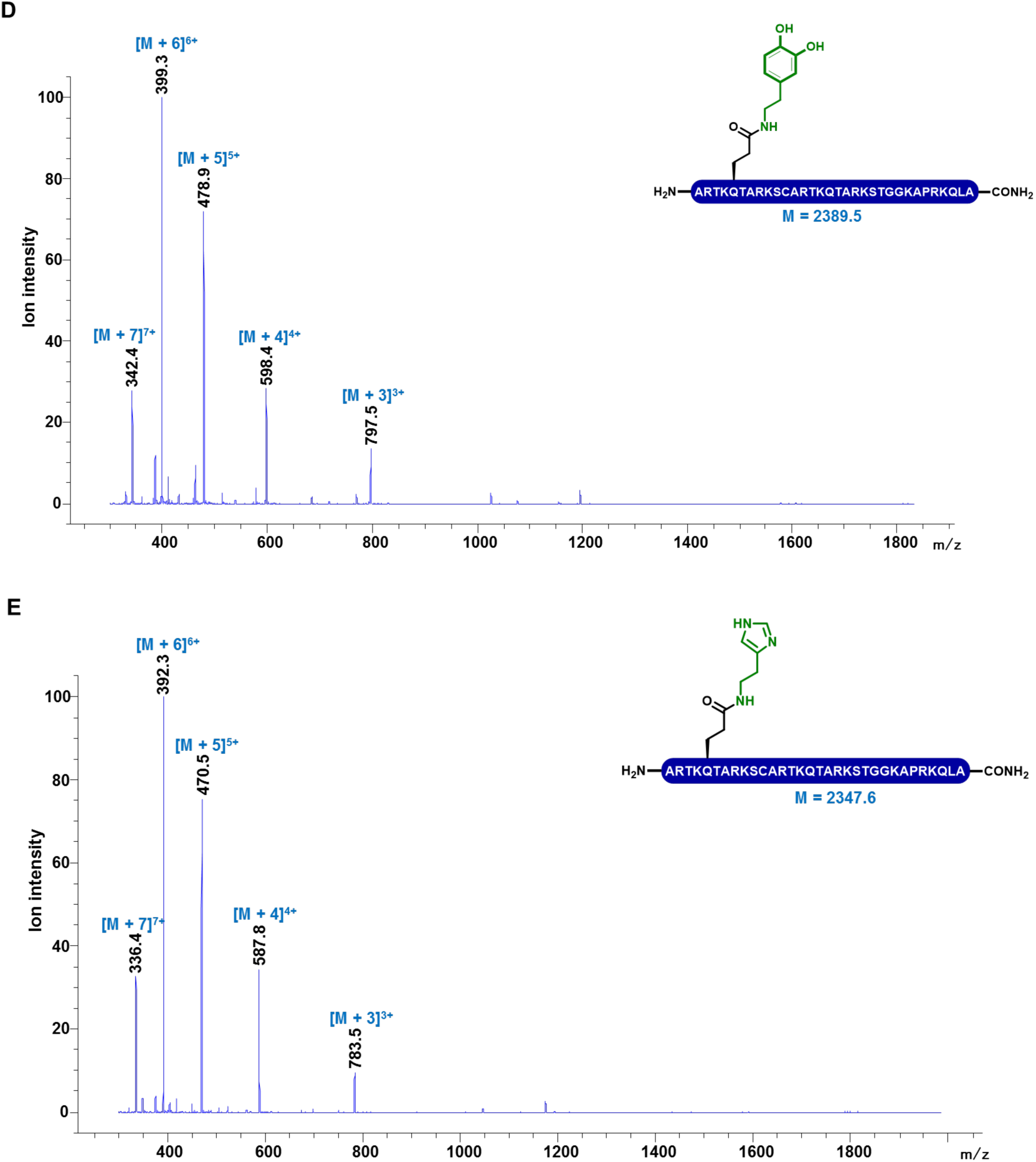
Structures and LC-MS analyses of the peptides utilized for *in vitro* biochemical reactions in this study. Side-chain protecting groups are omitted for clarity. The synthetic H3 N-terminal peptides (containing 21 amino acid residues) used as standards and substrates include: (A) H3Q5, (B) H3Q5E, (C) H3Q5ser, (D) H3Q5dop and (E) H3Q5his.

**Supplementary Figure 3.**
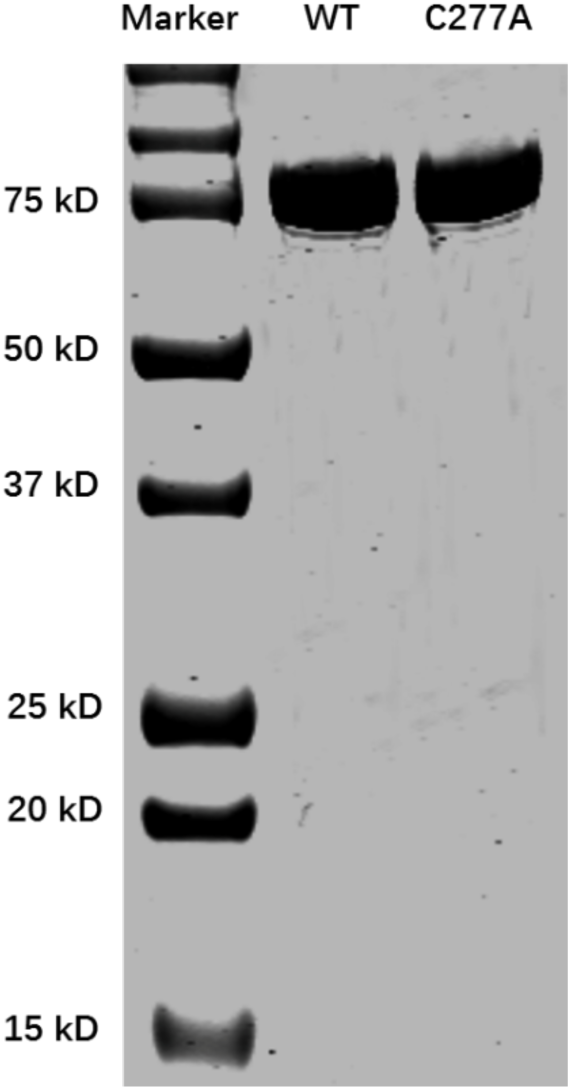
SDS-PAGE analysis of TGM2 purification. The lanes from left to right are protein marker, wild type TGM2, and TGM2-C277A mutant, respectively.

**Supplementary Figure 4.**
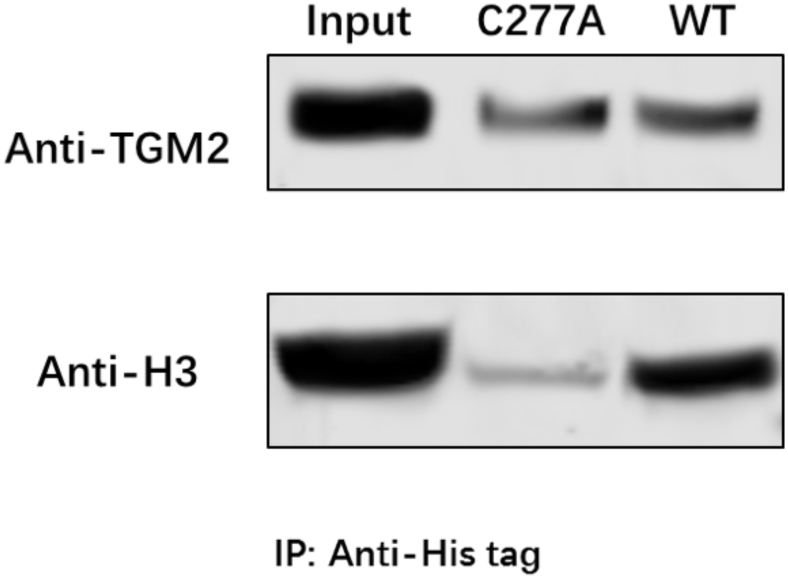
Identification of TGM2-H3 thioester complex from an *in vitro* biochemical reaction. 50 μM of free H3 proteins were treated with 50 μM of wild type TGM2 or TGM2-C277A mutant at 37 °C for 1 h. His8-tagged TGM2 was pulled down by BSA-blocked Ni^2+^-NTA agarose beads. The beads were washed, boiled, separated on SDS-PAGE and analyzed by western blot with anti-TGM2 and anti-H3 antibodies to detect the enrichment of H3.

**Supplementary Figure 5.**
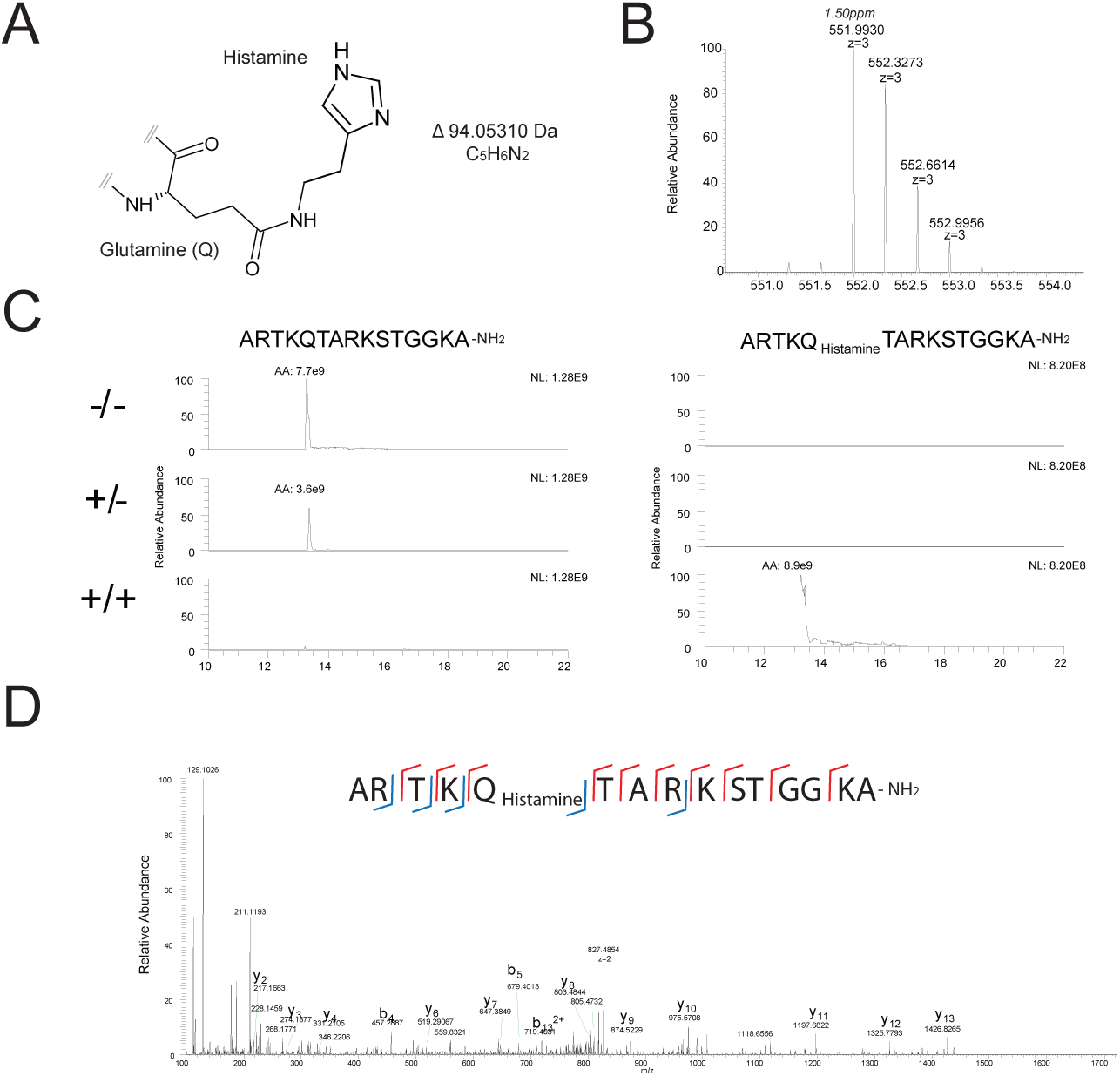
LC-MS/MS validation of H3Q5his on peptides. (A) Proposed structure of histaminylated glutamine. (B) High resolution/high mass accuracy mass spectrum of the triply charged histaminylated H3 tail peptide (ARTKQTARKSTGGKA-NH2; +histamine/+TGM2 condition). The difference between measured and expected mass of the amidated peptide was 1.5 ppm . (C) Extracted m/z (5 ppm) ion traces of the 2+, 3+ and 4+ amidated H3 tail peptide with (right panels) and without (left panels) histaminylated glutamine. Top, middle and bottom panels show signals measured under –/–, –/+ and +/+ (histamine/TGM2) conditions, respectively. Integrated area under curve are shown next to the peaks. Based on the extracted signals, the reaction is close to complete. (D) Tandem mass spectrum (35,000 resolution) of the doubly charged glutamine 5 histaminylated H3 tail peptide (+histamine/+TGM2 condition). Selected fragment ions (y and b) are labeled. Lowest mass was m/z 100. Vertical lines in red and blue within the peptide sequence are used to show matched peptide fragment ions.

**Supplementary Figure 6.**
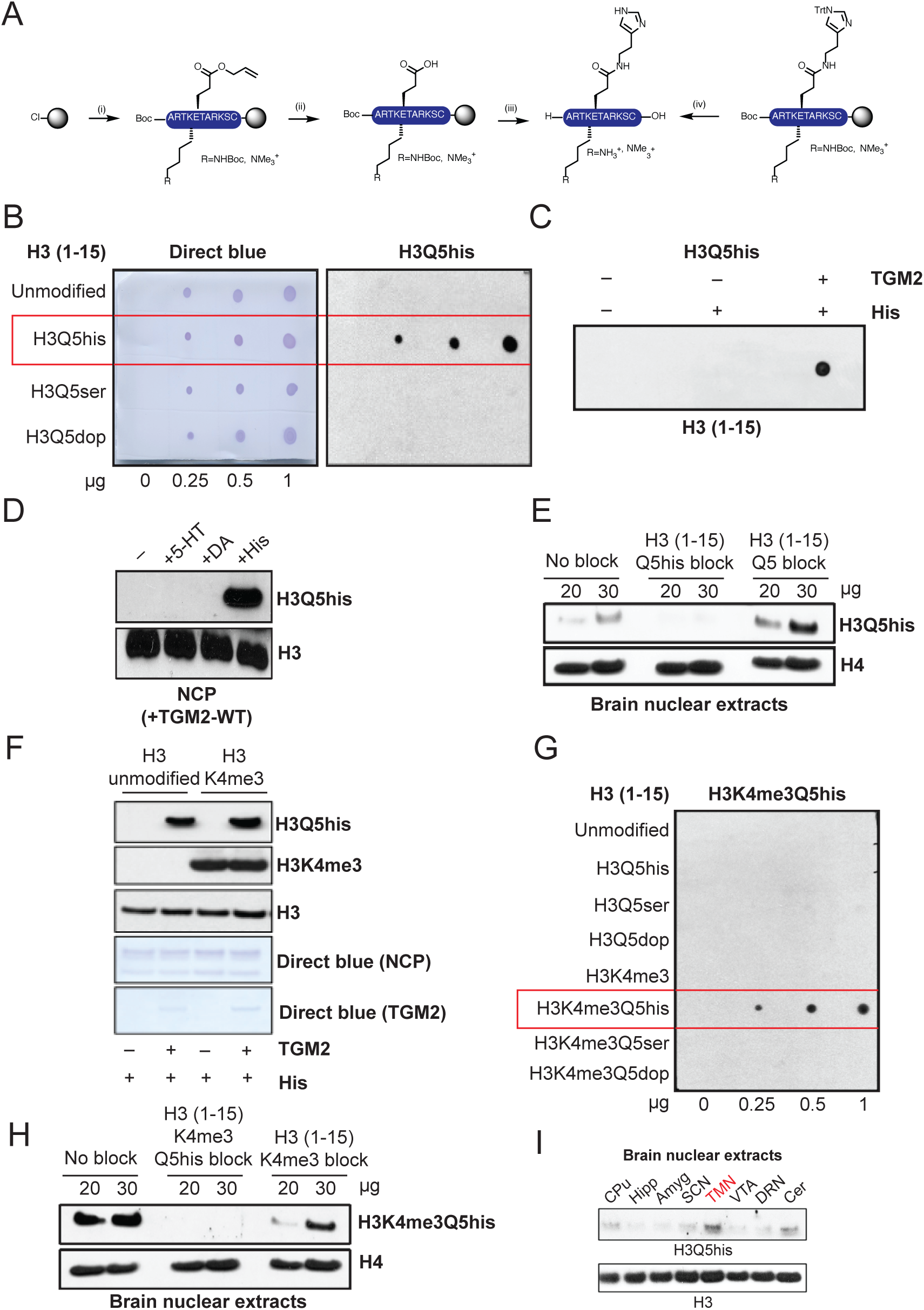
Generation and validation of H3Q5his antibodies. (A) Synthesis of peptide antigens on 2-Cl trityl resin by (i) iterative Fmoc solid-phase peptide synthesis incorporating Fmoc–Glu(OAII)-OH at position 5 and either Fmoc–Lys(Boc)-OH or Fmoc–Lys(Me_3_)-OH at position 4, (ii) followed by Pd(0) deallylation, (iii) coupling of Trt-protected histamine (iv) acidolytic cleavage from the resin and global deprotection. Side-chain protecting groups are omitted for clarity. (B) Peptide dot blot titrations testing the α -H3Q5his antibody’s specificity against unmodified *vs.* Q5his *vs.* Q5ser *vs.* Q5dop peptides. Direct blue staining was used to control for peptide loading. (C) Peptide dot blots testing the α -H3Q5his antibody’s reactivity -/+ histamine, -/+ TGM2. (D) Western blot analysis testing the α -H3Q5his antibody’s reactivity/specificity on NCPs following TGM2 mediated transamidation of histamine *vs.* serotonin *vs.* dopamine. Total H3 is provided as a loading control. (E) Peptide competition (i.e., no block *vs.* unmodified H3 block *vs.* histaminyl blocks) western blotting analysis of lysates from mouse brain indicating the specificity of our α - H3Q5his antibody. (F) Western blotting following TGM2 histaminylation reactions on unmodified *vs.* H3K4me3 mononucleosomes reveals that TGM2 can transamidate histamine to H3 in the context of adjacent H3K4me3. (G) Peptide dot blot titrations testing the α -H3K4me3Q5his antibody’s specificity against unmodified *vs.* Q5his *vs.* Q5ser *vs.* Q5dop *vs.* K4me3 *vs.* K4me3Q5his *vs.* K4me3Q5ser *vs.* K4me3Q5dop peptides. (H) Peptide competition (i.e., no block *vs.* H3K4me3 block *vs.* H3K4me3Q5his blocks) western blotting analysis of lysates from mouse brain indicating the specificity of our α -H3K4me3Q5his antibody. (I) Western blotting for H3Q5his across multiple brain regions in mouse reveals relative enrichment for the mark in TMN *vs.* other non-histaminergic monoaminergic and non-monoaminergic brain structures.

**Supplementary Figure 7.**
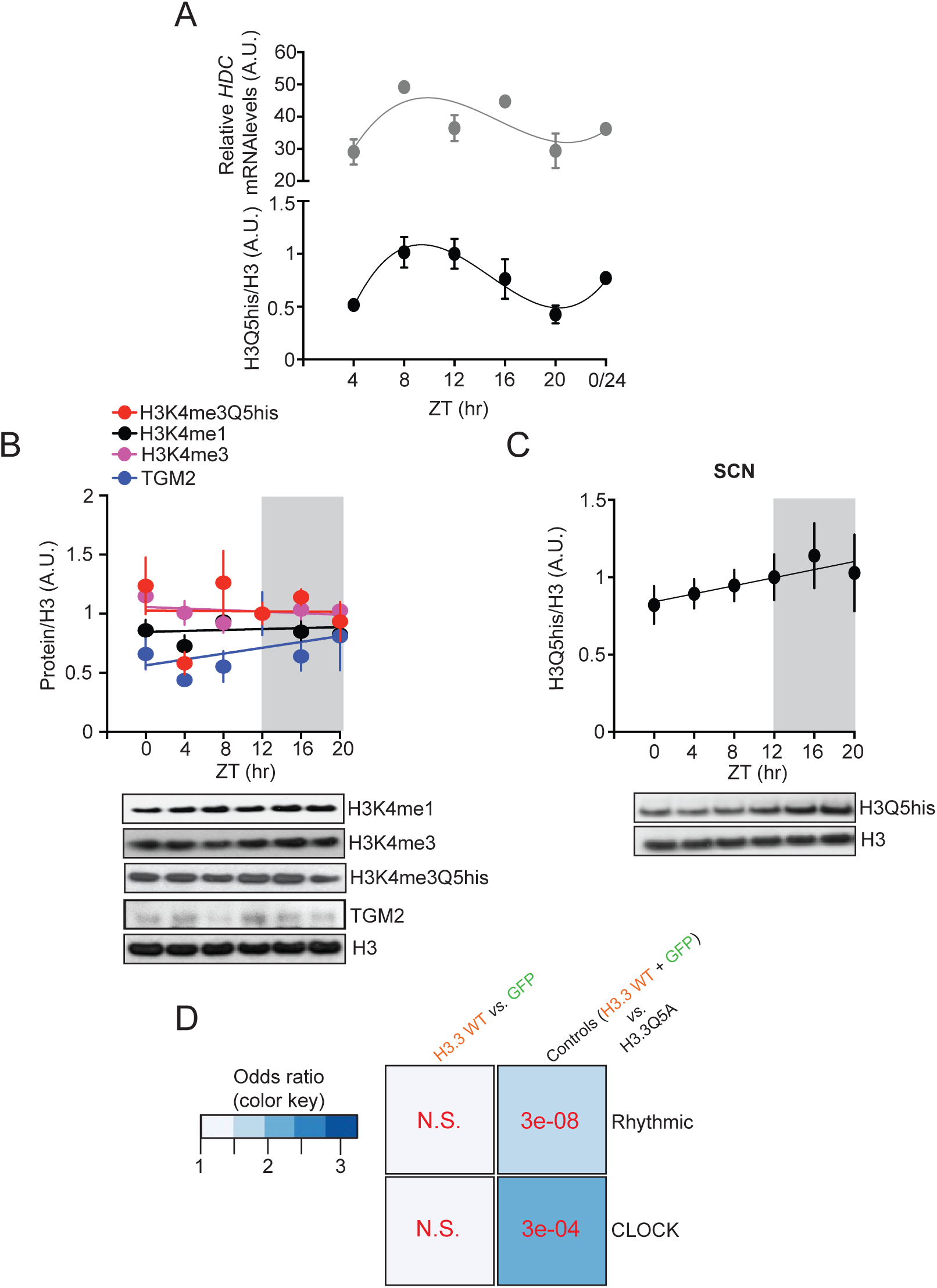
H3K4me3Q5his, H3K4me1, H3K4me3 and TGM2 do not display rhythmic expression in TMN, and H3Q5his is not rhythmic in SCN. (A) Comparison of H3Q5his/H3 expression (from Figure 4D) *vs. Hdc* mRNA expression (from RNA-seq presented in Figure 4A) in mouse TMN across the ZT revealed a similar pattern of expression, suggesting that alterations in H3Q5his may be dictated, at least in part, by local concentrations of Hdc, the enzyme responsible for histamine biosynthesis. (B) H3K4me1, H3K4me3, H3K4me3Q5his and TGM2 do not display rhythmic patterns of expression in mouse TMN across the ZT (all normalized to total H3). [*n*=7-10/time point; comparison of fits analysis between third order polynomial, cubic trend (alternative hypothesis), *vs.* first order polynomial, straight line (null hypothesis) – all p>0.05, null hypothesis accepted]. (C) H3Q5his does not display rhythmic patterns of expression in mouse SCN across the ZT (normalized to total H3). [*n*=8/time point; comparison of fits analysis between third order polynomial, cubic trend (alternative hypothesis), *vs.* first order polynomial, straight line (null hypothesis) – all p>0.05, null hypothesis accepted]. (D) Odds ratio analysis (Fisher’s exact test) comparing overlapping gene sets between RNA-seq data collected from H3.3Q5A *vs.* control (H3.3 WT + empty vector collapsed) transduced mouse TMN and rhythmic genes identified in Figure 4A, as well as ChEA identified CLOCK gene targets. p-values are provided. Data are represented as mean ± SEM.

**Supplementary Figure 8.**
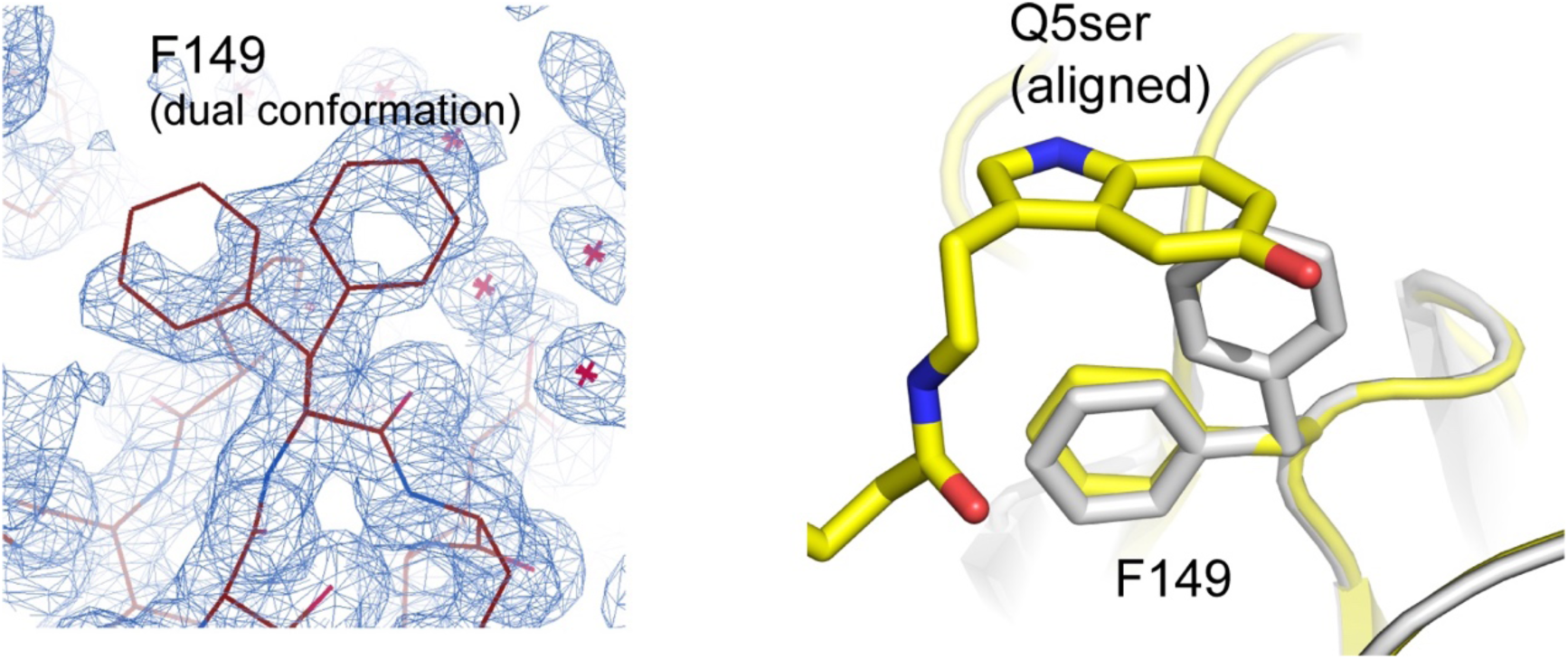
H3Q5ser flexibility in the H3Q5ser-WDR5_WD40_ structure. Electron density of the WDR5-F149 residue shown in COOT. F149 displays a double conformation in the structure. H3Q5ser is aligned here based upon the WDR5-H3Q5ser structure (PDB ID: 7CFP).

## Notes

### Competing Interest Statement

The authors have declared no competing interest.

### Summary of Updates

Addition of co-author, Lauren Dierdorff

